# Structural network comparison of domains across remote homologues arising from different domain architectures

**DOI:** 10.64898/2025.12.19.695395

**Authors:** Sidhanta Swayam Prakash Das, Vasam Manjveekar Prabantu, R. Sowdhamini, N. Srinivasan

## Abstract

The homology of protein domains indicates an evolutionary relationship between them. Remote homologues, on the other hand, share very low sequence similarity while maintaining a highly similar structural topology. These similar structures can be compared using several metrices, but a comparison of networks built from amino acid residues to mimic non-covalent interactions provides a better understanding of their similarities or minute differences. Comparison of structurally related domains are studied to some extent, while the impact of domain combinations is poorly studied. In this study, several types of pairs of domains, arising from different multi-domain proteins were compared to understand the effect of cognate domain on the structural architecture of remote homologues. Several network parameters were compared among these domains. We found that an interacting domain of unrelated structure exerts higher structural and evolutionary pressure on the remote homologues, leading to structural embellishments and functional modifications. This exerted change is mediated by local rearrangements of the amino acid residues in the structure. Case studies were conducted to understand these effects. These results will assist in re-defining the homology among domains, which would rather help in better functional annotations and structural modelling.

## 1. Introduction

The functioning of proteins is attributed to protein domains, which are otherwise called as structural, functional and evolutionary units of proteins. They are often known as building blocks since their presence, interactions, and orientations confers distinct biological roles (Basu et al., 2008, 2009). The arrangement of these domains has been the subject of attention in the past, mainly focusing on the understanding of the evolutionary history of proteins they constitute (Chothia et al., 2003; Jin et al., 2009; Margelevičius & Venclovas, 2010), assessment of homology detection (Chakrabarti & Sowdhamini, 2004a), fold recognition approaches (McGuffin et al., 2001), functional annotations (Bonello & Orengo, 2024) and many more. On the other hand, there are several unrelated proteins with diverse sequences and functions, whose structures are very similar as the structural folds are limited in nature (Chothia, 1992). This also implies that structure is more conserved than sequence (Illergård et al., 2009). There is unequal distribution of sequences to adopt folds (Iyer et al., 2018) showing higher preference for certain structural classes. Some structural embellishments could be observed in the common core (Cuff et al., 2008; Reeves et al., 2006), including additional secondary structures (Sandhya et al., 2009) which may provide support for evolving new functions. Such domains are organised into a database for easier study of superfamilies (Mutt et al., 2014).

In addition, the functional diversity of proteins using a limited repertoire of domains is achieved by several domain combinations in a single polypeptide chain to form a multi-domain protein. Therefore, a majority of the proteins in organisms are multi-domain in nature (Gerstein, 1998; Teichmann et al., 1998). The constituent domains in multi-domain proteins are mostly duplicated rather than adopting new domain architectures (Apic et al., 2003). Wherever, there exists a multi-domain protein with different domain superfamily combinations, it is mostly observed that the sequential order of the occurrence of these domains is preserved, since these combinations are thought to be derived from duplications of domain combinations that arose first as a result of single recombination event (Bashton & Chothia, 2002; Vogel et al., 2005). The extent of the duplication of these domain combinations depends on natural selection (Vogel et al., 2005) and therefore multi-domain proteins contain a limited set of domain superfamily pairs. This natural selection prefers proteins of certain fold combinations for geometric advantages during folding and to lower the optimization cost for function of multi-domain proteins (Naveenkumar, Kumar, et al., 2019). Overall, these domain duplications in multi-domain proteins increases the abundance, while new recombination increases the versatility of proteins (Vogel et al., 2005), which is analogous to the fact that protein universe is wide but limited in terms of number and uniqueness of domains (Nacher et al., 2010). Moreover, the functions of proteins of identical domain combination (Hegyi & Gerstein, 2001) and their global dynamics (Keskin et al., 2000) are much similar. Hence, fine-tuned functional assignments are usually preserved and can be derived for proteins on the basis of identical domain architectures.

However, minute structural differences among homologous domains, need to be analysed which enables altered and non-identical protein function. For such structural comparisons, several well-known metrices are used in literature, including global distance test total structure (GDT-TS), TM-score and root mean square deviations (RMSD). But these metrices mostly consider the structural backbone to compare them. With the recent advances in network biology, which considers the macromolecule as a single connected entity, it is possible to differentiate minute differences in both local and global level. This Network Similarity Score (NSS) considers non-covalent interactions stored in matrix form and uses its spectral features to identify sources of differences (Gadiyaram et al., 2017, 2019). In recent works, NSS was proven to be excellent scoring scheme to elucidate structural features distinguishing SARS-CoV2 spike protein conformers (Halder et al., 2020), proteins of transient interactions (Prabantu, Tandon, et al., 2023), disease causing mutations from wild type variant (Prabantu et al., 2021) and so on. Such comparisons also help to understand the evolution and functional similarities of homologous proteins (Prabantu, Gadiyaram, et al., 2023), to understand the stabilising factors of TIM barrel structure despite high sequence dissimilarity (Vijayabaskar & Vishveshwara, 2012), to understand the structural details of Rossmann fold domains, co-existing with catalytic superfamily domains (Bashton & Chothia, 2002), to uncover Pleckstrin homology (PH) domain specific features (Naveenkumar, Sowdhamini, et al., 2019), and so forth. Many in depth studies focusing on homologous domain comparisons and effect of domain architectures are also available (Bashton & Chothia, 2007; Chakrabarti & Sowdhamini, 2004b; Dessailly et al., 2010; Itoh et al., 2007; Lees et al., 2016; Roy et al., 2009; Thangudu et al., 2008; Todd et al., 2001; Vogel, Berzuini, et al., 2004). However, structural comparisons of homologous domains, which provide better understanding of the relationship of structure, function and evolution are limited. Domains from the same superfamily were compared to analyse divergences of structural topology within superfamilies. These comparisons were accompanied by different types of domain comparisons arising from different domain combinations in multi-domain proteins. Differences among domain comparison types were observed in terms of their degree variability, conservation of hubs and finally the network dissimilarities. Although domains are compared from the same superfamily in CATH, domains of proteins with same domain architecture showed least network variability in both local and global levels, while domain comparisons from two different domain architectures showed the maximum variability. Interestingly, it also observed that domains within a single chain forming structural repeats showed some variability, despite domain superfamily duplications in the evolution. This peculiarity is mostly due to domination of a single domain to perform the function, as inferred from case studies. These results will help to redefine the definition of homology among structural domains, which will greatly impact protein structure and function predictions.

## 2. Results and discussions

The initial dataset consists of monomeric multidomain proteins, where the number of interacting domains range from two to eight (Supplementary Figure S1). The number of multi-domain proteins decreases as the number of constituent domains increases. This agrees with the fact that majority of the multi-domain proteins consists of at least two domains, and the shorter domain combinations are reused for easy functional modification (Apic et al., 2003; Vogel, Bashton, et al., 2004). The number of domains in a protein follows power law distribution (Nacher et al., 2010), which is also observed in our dataset consisting of 594 two-domain proteins and only three proteins with eight domains (Supplementary Figure S1). Therefore, to mimic the properties of multidomain proteins, we used two-domain proteins to perform our analyses (similar to (Bashton & Chothia, 2002)), which in turn will also eliminate the interference effect by other cognate domains. These two-domain proteins can be seen as simple representations of higher order multi-domain proteins. We used NOXclass to predict domain interactions as permanent and transient interactions, but this concept is not used in the study. However, this classification prediction strengthens our dataset by adding additional filters to select variety of interacting domains (please see Methods for details). Moreover, previous studies have shown that mutations bring changes to protein folding (Lorch et al., 1999; Taverna & Goldstein, 2002; Tokuriki & Tawfik, 2009) and stability (Lorch et al., 2000) of protein, dynamics of some intrinsically disordered proteins (Seera & Nagarajaram, 2021) and residue network of the proteins (Prabantu et al., 2021). To avoid such bias due to mutations, we removed proteins with engineered mutations by taking help of filters in RCSB PDB (Berman et al., 2000b).

We used protein structural networks (PSNs) as mentioned in the methods, to enumerate the number of atomic contacts per kind of residue type (Supplementary Figure S2). We observed that Histidine to Tryptophan interacts with each other using the largest number of atomic contacts (50 atomic contacts), followed by Tryptophan-Tryptophan (48 atomic contacts) and Tyrosine-Histidine interactions (48 atomic contacts). This observation holds true as these large atomic contacts arise from larger ring-containing amino acids as shown in Supplementary Figure S3. On the other hand, the lowest number of atomic contacts arise mostly from backbones of Glycine-Glycine interactions.

Using these proteins in the dataset, we grouped the domains based on the CATH IDs till their ‘H’ group (homologue) implying that they have a common ancestor with shared structural and functional similarities among the domains. Figure 1 illustrates the distribution of domain superfamilies in our dataset. Our dataset contains several singleton homologous superfamilies containing a single domain, while a few containing many domains, following the previously known uneven or power-law distribution of domain families (Koonin et al., 2002; Qian et al., 2001). Trypsin-like serine proteases (CATH: 2.40.10.10), Periplasmic binding protein-like II (CATH: 3.40.190.10), Response regulator (CATH: 3.40.50.2300), etc. are some of the highly occurring domain superfamilies in our dataset, as well as in the CATH database (Orengo et al., 1997). The large duplications of a few domains make the protein universe wide but thin (Nacher et al., 2010).

**Figure 1.**
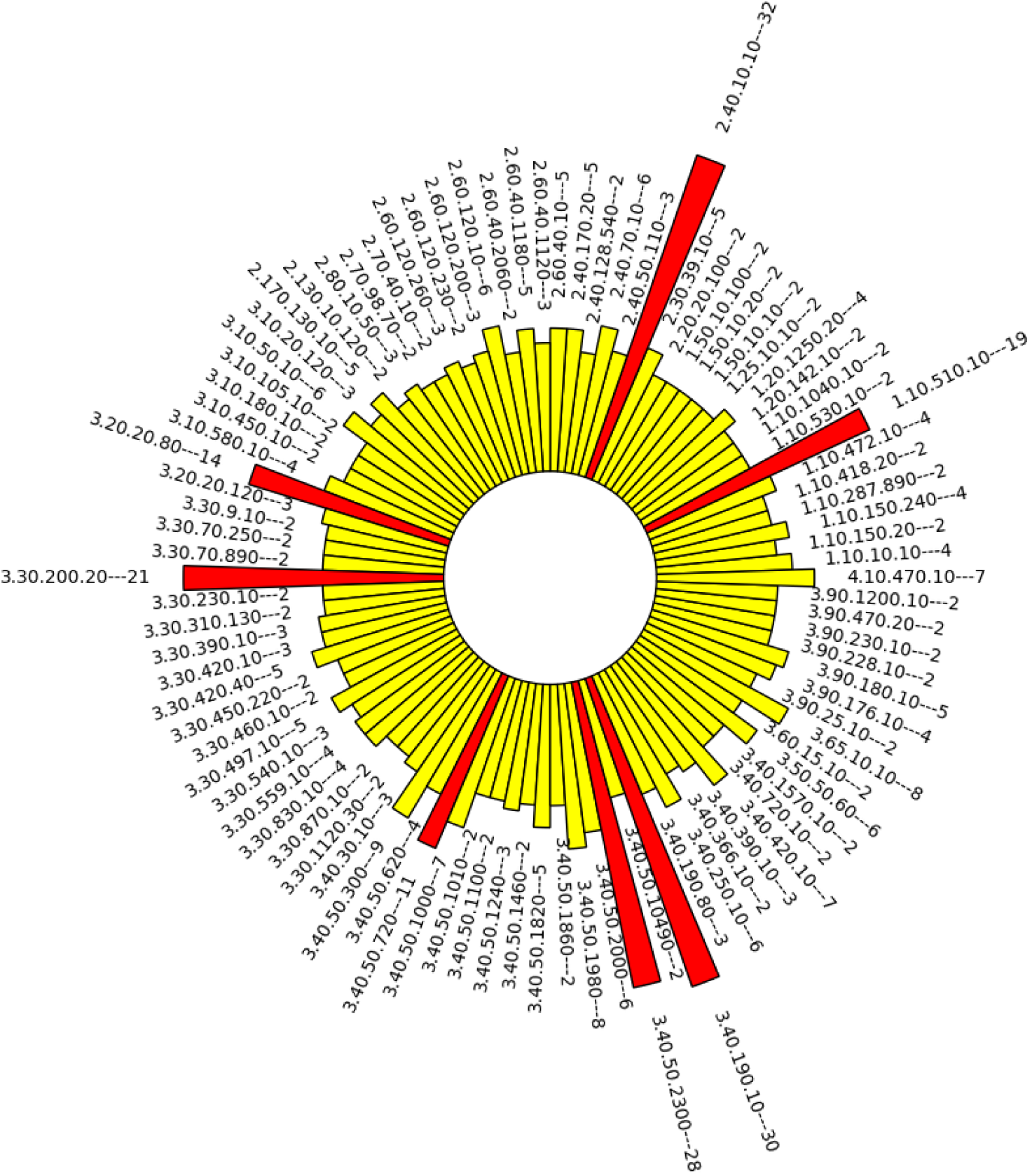
Domain frequencies in the dataset. The bars represent a CATH homologous superfamily, and each bar is labelled with its CATH id as well as the count of members in the group from the dataset, separated by a delimiter. The bars are coloured red if the membership is more than 10.

From the dataset of interacting two domain proteins to represent multi-domain proteins, we created subsets consisting of pairs of domains based on the domain neighbourhood type. The first type of domain comparisons includes pairs of interacting domains of remote homology in the same polypeptide chain (we refer this as “Type 1”). The second type (referred as “Type 2”) includes pairs of domains from the same homologous superfamily, but from the two different proteins where their corresponding interacting domains are also from the same superfamily. The third type (“Type 3”) includes pairs of domains from the same homologous superfamily from two different proteins, where their corresponding interacting domains do not share any structural evolutionary history. A graphical representation of the types of domain comparison is presented in Figure 2 (shown for simplicity), while all possible combinations used in this study are shown in Supplementary Figure S4. These domains from the same superfamily are thought to share common ancestry. However, the local environment of the domain with respect to the interactions arising from their partners differs in each type. These types will highlight the differences in the PSNs of domains within a multi-domain architecture

**Figure 2.**
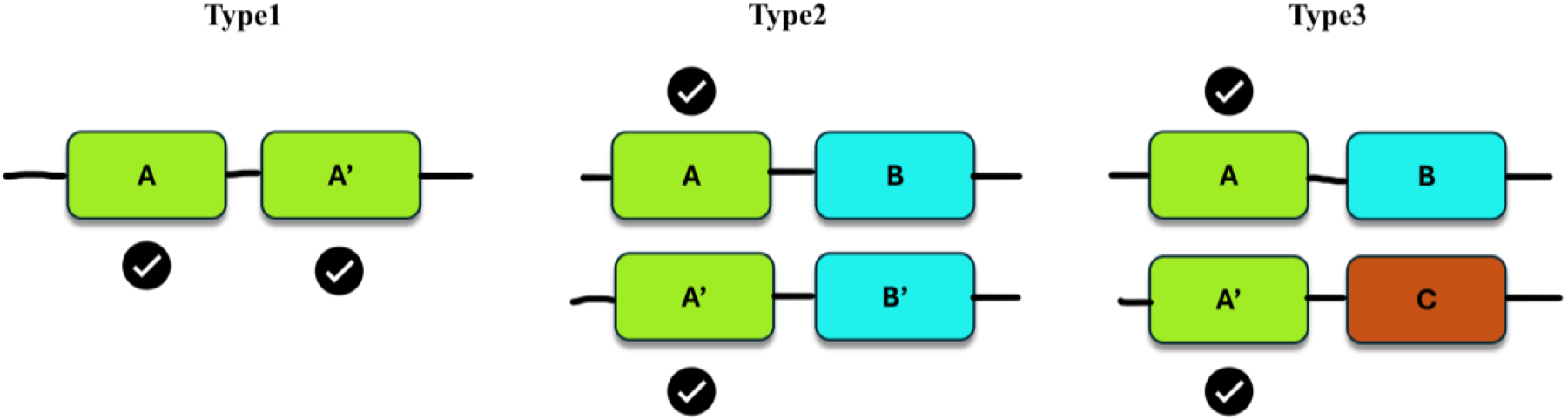
Different types of domain pairs in the study. The homologues are marked with ‘(single apostrophe) mark and colours. The domain pairs to be compared are marked with a tick mark.

(Type1) versus domains from same multi-domain architecture (Type2) versus domains from two different multi-domain architectures (Type3). The number of pairs resulting for each type with these architectural combinations make them unequal, where type1, type2 and type3 have 46, 834 and 270 pairs respectively. This asymmetry is mostly due to the evolutionary natural selection of certain domain combinations in nature based on their importance (Vogel et al., 2005). The current dataset also includes domain comparisons arising from two proteins, where one of the proteins has Type1-like architecture, while maintaining the rule of domain comparison in different types. These kinds of pairs are called peculiar pairs in this study. For example, in the comparison of N-terminal domains in 1DUA and 1P3C, both proteins have Type1 like architecture, while following the rule of Type 2 domain comparisons. The number of such peculiar pairs in Type 2 and Type 3 are listed in Supplementary Table S1.

From the preliminary analysis (as detailed in Supplementary Figure S5) on these pairs from different types of comparison, we observed different ranges of domain length differences. Type 1 and Type2 pairs have a maximum domain length difference of around 100 (with a few outliers), while Type3 have some pairs with more than 100 length difference (comparatively higher tail height in the histogram). Length variations among some domains in the superfamilies are associated with length-deviant and length-rigid superfamilies, rather than an additional domain in the protein chain (Mutt et al., 2014; Sandhya et al., 2009). When the sequences of the domains were compared, most domain pairs retain a maximum sequence identity of 40% (few in the 40-50% range). These sequence identities go as low as ∼10%, where many pairs are in lower ranges, suggesting their remote homology within the same homologous superfamily (Supplementary Figure S5 B, S5 D, S5 F). Type1 has a domain pair which is a clear case of tandem repeat (Björklund et al., 2006; Kajava, 2012), with very high sequence identity (PDB ID: 3KA2, where domains have 99% sequence identity and same hierarchy till CATHSOLI). All types of comparisons have a good topological equivalence after removing poorly aligned structures (TM score<0.5). Moreover, different domain pairs retain similar extents of sequence changes, while accommodating some structural changes, possibly to accommodate new divergent functions. These vertical lines can also be noted as divergent sequence subgroups. Hence, this dataset covers domain pairs having duplications as well as convergent/divergent evolutions. A higher sequence divergence with lower structural divergence also highlights the fact that structural domains are limited and more conserved than sequences (Illergård et al., 2009), so re-used often in the protein evolution.

### 2.1. Domains from two different domain combinations have higher degree variability

In a PSN, which is a node-edge representation of amino acids in the domain, a degree of a node is the number of interactions in the form of edges, connecting to its interacting partners. The degree parameter of a node in the network suggests how well the node is connected in the network implying its importance in the network. Using the PSN, we calculated the degree of each node in the domain structural network and compared the topological equivalent nodes for its degree with the other remote homologous domain in the pair. We found that most of the nodes in the dataset have a degree of six and a highest degree of 19 (Supplementary Figure S6) arising from a Tryptophan residue. Although domains from different domain combinations are used, their degree distribution is very similar. The degree distribution in this study looks like a random network as the distribution somewhat follows a Poisson distribution rather than power law (Albert & Barabási, 2002). The power law like degree distribution is not a universal characteristic of real networks, and it deviates from this when the network is highly densely connected (Scholz, 2010). Protein domains are highly connected entities which could be the reason behind such dense network to give such a distribution. Further, we calculated the degree differences for the equivalent residue pairs and calculated variances for each pair of domains for degree difference variability, which would imply the degree variability of pair of domains from same superfamily affected by domain interactions.

From Figure 3A, it is evident that domain pairs from Type3 have a comparatively higher degree variability than Type1 and Type2. It is also observed that, Type2 pairs have marginally lower degree variability than Type1, making them the pairs with lowest degree variability. The means of the degree difference variances can be found in Table 1, which quantifies the above observations. We also compared this degree variability with the Root Mean Square Deviations (RMSD) to understand the relationship between residue interactions in local environments and global structure. We found that, all domain comparison types have similar trend of increasing RMSD along with increase in degree variability (Figure 3B), with observable differences when a linear regression line is drawn. Type1 domain pairs show some correlation as inferred from the plot, but a linear relationship is hard to establish from correlation coefficients (Table 1). Type3 domain pairs have a higher distribution of RMSD values along with higher degree differences. This, in turn makes the relationship between these structural parameters stronger (spearman correlation coefficient of 0.702) than other two types of domain comparisons (Type1 and Type2). It is also noted that the spearman’s correlation values of Type2 and Type3 have minute differences (Table 1), suggesting that the strength and direction of the relationship between degree variability and RMSD are similar. However, interpretation of linear regression lines implies that Type3 domain pairs show higher RMSD increase than Type2 pairs for similar pairwise degree difference variance. We also compared these degree difference variances to another structural comparison metric, TM-score, and we observed a negative correlation, while Type3 pairs had consistently lower TM-scores, giving a worse correlation (Supplementary Figure S7). When a pair of remote homologous domains from Type 1 and Type 2 are considered, it is observed that higher degree difference variance is associated with improper domain superposition which are due to comparatively lower structural similarity. These observations implies that remote homologous domain pairs from Type3 have larger differences in the local residue interactions. This can lead to larger differences in the flexibility of the local regions around the residues, thereby easier propagation of this local phenomenon to the global backbone deviations, and vice-versa. On the other hand, Type1 and Type2 local interaction variations are similar, either due to their interactions in a single chain or related structural domain combinations, which are the possible reasons for maintaining a similar local interaction, hence, smaller deviations in their degrees.

**Figure 3.**
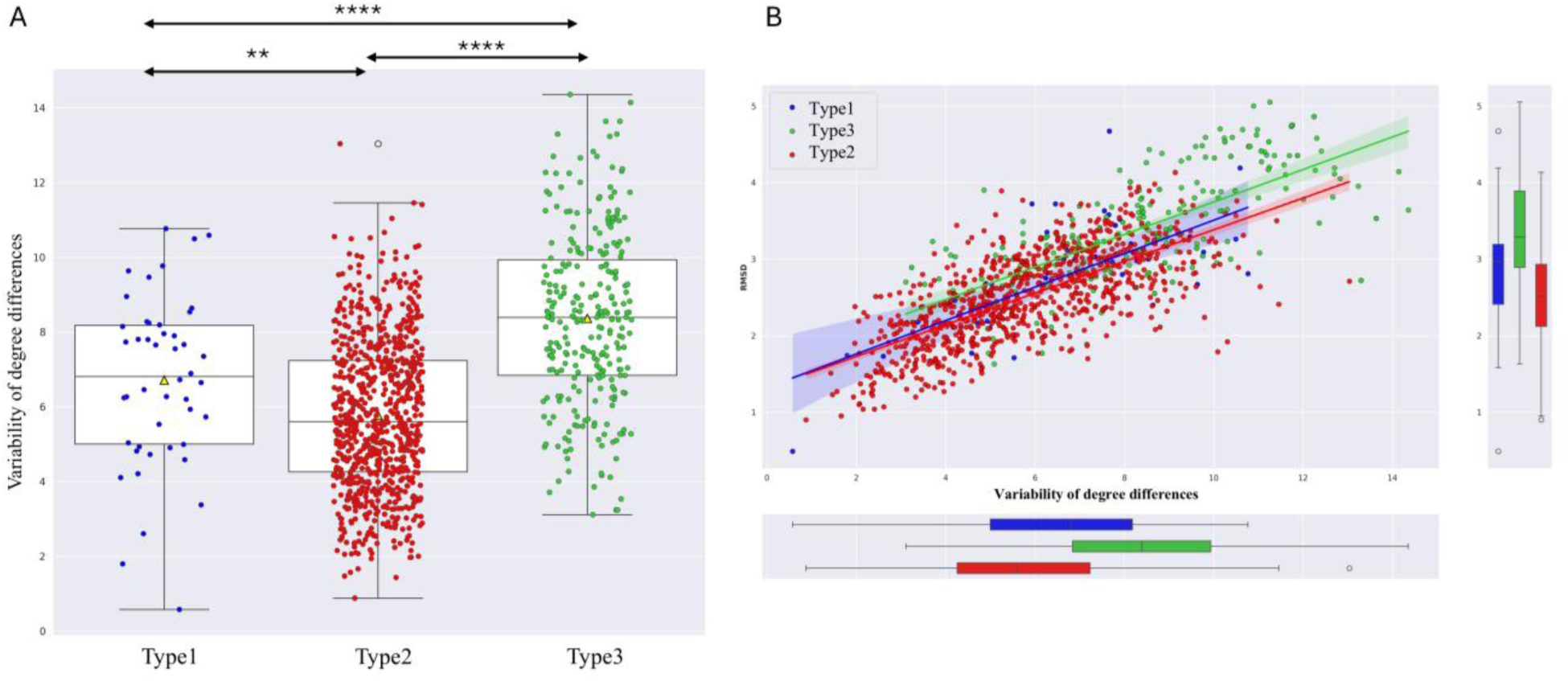
Degree variability among domain comparison types. **A.** The variances of topological degree differences of each domain pair. The Mann Whitney U p-Values are 0.001, 2.9*10^-49^, 2.6*10^-05^ for Type1 vs Type2, Type2 vs Type3 and Type1 vs Type3 respectively. B. The relation between degree difference variances (degree variability) to structural backbone difference (RMSD). The distribution of the variables is plotted in the marginal plots and linear regression lines are drawn in the scatterplot.

**Table 1.** Statistics of Degree variability. Mean of degree variability across domain comparison types and their correlation with RMSD.

| Types | Mean of degree variances | Spearman correlation coefficient |
| --- | --- | --- |
| Type1 | 6.708 | 0.599 |
| Type2 | 5.775 | 0.699 |
| Type3 | 8.434 | 0.702 |

### 2.2. Distinct hub distribution of domain pairs based on differences in combination of domain interactions

The amino acid residues with the highest degree, implying highest number of interactions with other amino acids are known as hubs in the network. These hubs have multiple connections in the network, through which many other residues in the network also interact with other residues, hence maintaining the network. Like other real networks, these structural networks are also susceptible to targeted attack on hubs (Cohen & Barabási, 2002). Therefore, hubs are also known as hotspots of networks.

Here, we calculated the degree of all the residues in the dataset, and their distribution was plotted (Supplementary Figure S6). We observed that a residue with degree 11 is above the rest 90% of the degrees (90 percentile). Hence, we used a degree cutoff of 11 to define a residue as hub in the network. We counted the number of hubs in each domain in the dataset and later, defined the topological residue pairs from the domain comparisons as conserved hub or non-conserved hub based on their preservation of hub status. It was observed that all the domain comparison types have a larger number of non-conserved hubs than conserved hubs (Figure 4). This is similar to observations from few previous studies, where net loss and gain of hubs is higher than invariable hubs (Prabantu et al., 2021; Prabantu, Tandon, et al., 2023). This consistency suggests a distinct way of maintaining structural integrity across domains from the same superfamily, despite maintaining highly similar structures. However, differences arise when we consider different pairs of domain comparisons (Types). When a linear regression line was drawn to infer the relation between the number of conserved and non-conserved hubs, we found that Type3 members have the best spearman correlation coefficient pointing towards the best linear relationship and this line describes a better linear relationship than Type1 and Type2. We also calculated the slope of this line, to compare the rate of change of each hub type, viz, conserved and non-conserved hubs. A slope of 2.23 (Table 2) was obtained for Type3 which suggests that rate of change of non-conserved hubs is more than two times the number of conserved hubs when the domain pairs are compared. On the other hand, Type2 domain pairs showed slope near to one, marking this type as the least rate of hub change. This suggests the rate of change of conserved and non-conserved hubs are similar as the number of hubs increases in the domains of Type2. It is to be noted that Type3 pairs form two minor clusters in the plot, where these clustering patterns are due to alignment lengths between domain pairs (Supplementary Figure S8), therefore, higher hubs. We also found that the distribution of hubs in domain comparison types (Type1 to 3) varies only due to distribution of non-conserved hubs while the preservation of some hub patterns is similar (Supplementary Figure S9). This suggests that among remote homologous domains with different domain interactions, the structure is maintained by some of these conserved hubs, while non-conserved hubs are responsible for the structural embellishments for their selected function due to evolutionary pressure/selection. These observations point to a conclusion that although remote homologues maintain their overall structure, their residue interaction varies from types of domain combinations. The overall structure is preserved by a few hubs, while many hubs are located strategically to prioritize the evolution needed for different domain combinations to promote diverse function.

**Figure 4.**
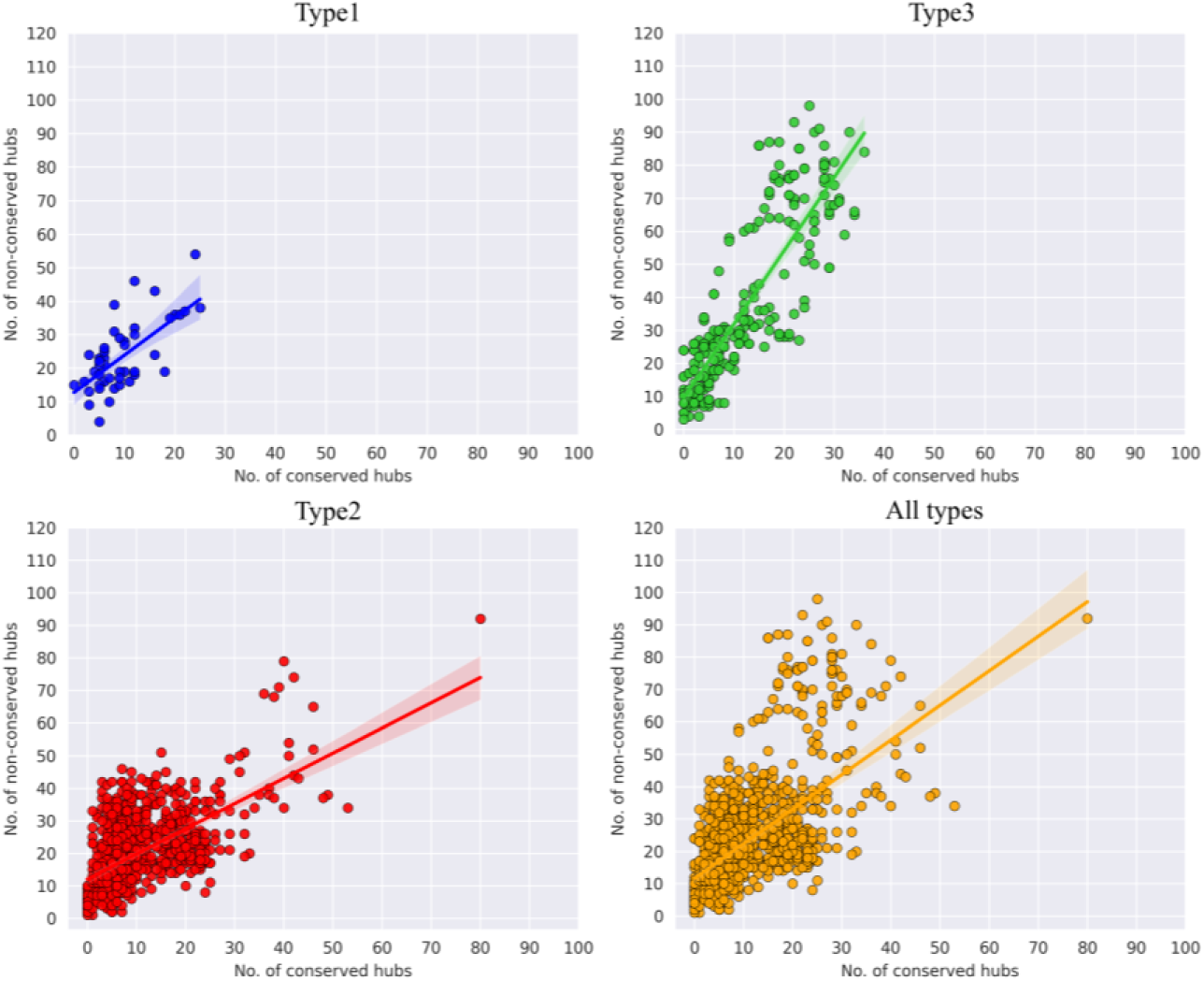
Hub preservation in different domain comparisons and all pairs. The lines through the scatter plot show a linear regression line.

**Table 2.** Linear correlation between conserved and non-conserved hubs. The rate of change non-conserved hubs with respect to conserved hubs can be inferred from the linear equation slope.

| Types | Spearman Correlation Coefficient | Linear regression equation |
| --- | --- | --- |
| Type1 | 0.618 | $Y=1.75X+12.516$ |
| Type2 | 0.588 | $Y=0.78X+12.455$ |
| Type3 | 0.862 | $Y=2.23X+10.369$ |

### 2.3. Global network variability among remote homologues differs based on domain-domain interactions

Residue networks of two domains from the same superfamily were compared in both local and global levels using the Network Dissimilarity Score (NDS). This method aligns the spectra of the network matrix to ensure the best alignment of eigen vectors. NDS scores consist of its components which not only contribute to the final score but also give information at the local and global levels in a network (Gadiyaram et al., 2019). NDS takes a value from 0.08 to 0.487 in the current comparisons.

We compared the NDS of the domain comparison types to examine the differences among different domain comparisons arising from different domain combinations (Figure 5A). We observed a significant difference among three types of comparisons. Type 2 pairs showed the lowest mean of 0.29 and standard deviation of 0.056, while Type 3 pairs showed the highest mean of 0.357 and standard deviation of 0.058. These values were compared with single domain monomeric proteins from previous studies. We observed that conformers of a single domain protein (Prabantu et al., 2022) retain the lowest network dissimilarity ranges (with a mean of 0.113±0.04), followed by single domain homologous proteins from the same family (mean NDS= 0.249) (Prabantu, Gadiyaram, et al., 2023). It is noteworthy that the network dissimilarities of distant homologous from a superfamily, with different conjoint interacting domains, have comparatively higher network dissimilarities than close homologous in a family. This trend can be consolidated due to the type of relationships of domains and their interacting domains. The relationships, in turn, are largely due to the evolutionary pressure for maintaining structure and diverse functions.

**Figure 5.**
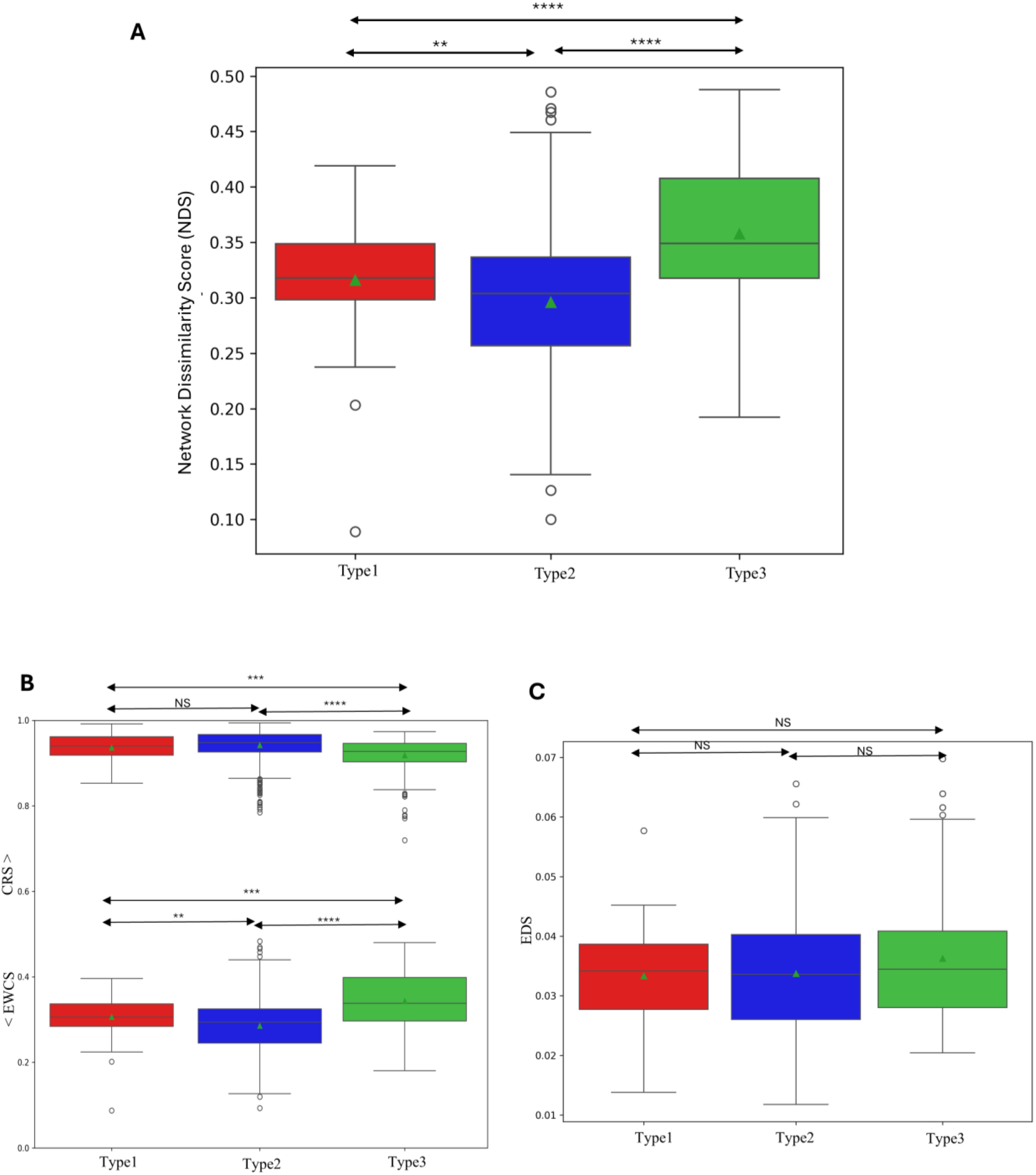
Network dissimilarity scores and its components. **A.** Distribution of network dissimilarity scores among types of remote homologues. Components of the NDS like EWCS and CRS are shown in B, and EDS in C. The mean of the distributions are shown as a green triangle within the boxes. The MannWhitney U p-Values of NDS are 0.0095, 3.07*10^-40^ and 4.59*10^-5^ for Type1 vs Type2, Type2 vs Type3, and Type1 vs Type3 respectively. P-values for CRS of Type1 vs Type2, Type2 vs Type3, and Type1 vs Type3 are 0.07, 1.02*10^-23^, and 0.001 respectively. For EWCS, 5.5*10^-3^, 6.097*10^-33^,0.2*10^-3^ in the same order.

From previous studies, it was observed that the network variability is more sensitive than backbone C^α^ RMSD implying compensation of structural similarity by small network re-arrangements in closely related domains (Prabantu et al., 2022; Prabantu, Gadiyaram, et al., 2023; Prabantu, Tandon, et al., 2023). We also compared the network dissimilarities with backbone RMSD to know if the network of inter-atomic interactions propagates to global structural deviations. From Figure 6, we observed that all the domain comparison types have some amount of linear relationship inferred from the spearman correlation (Table 3). Network dissimilarities and RMSD of Type1 and Type2 do not properly fit a linear relationship and there exist an asymmetric relationship. This asymmetric relationship suggests that remote homologues when interacting in a single chain or interacting with similar neighbour domains in a chain, largely re-orient their side chain and main chain interactions to minimally alter the global backbone trace. On the other hand, the side chain and mainchain interactions are equally reciprocated to backbone orientations very well for Type3 pairs involving two different domain combinations.

**Figure 6.**
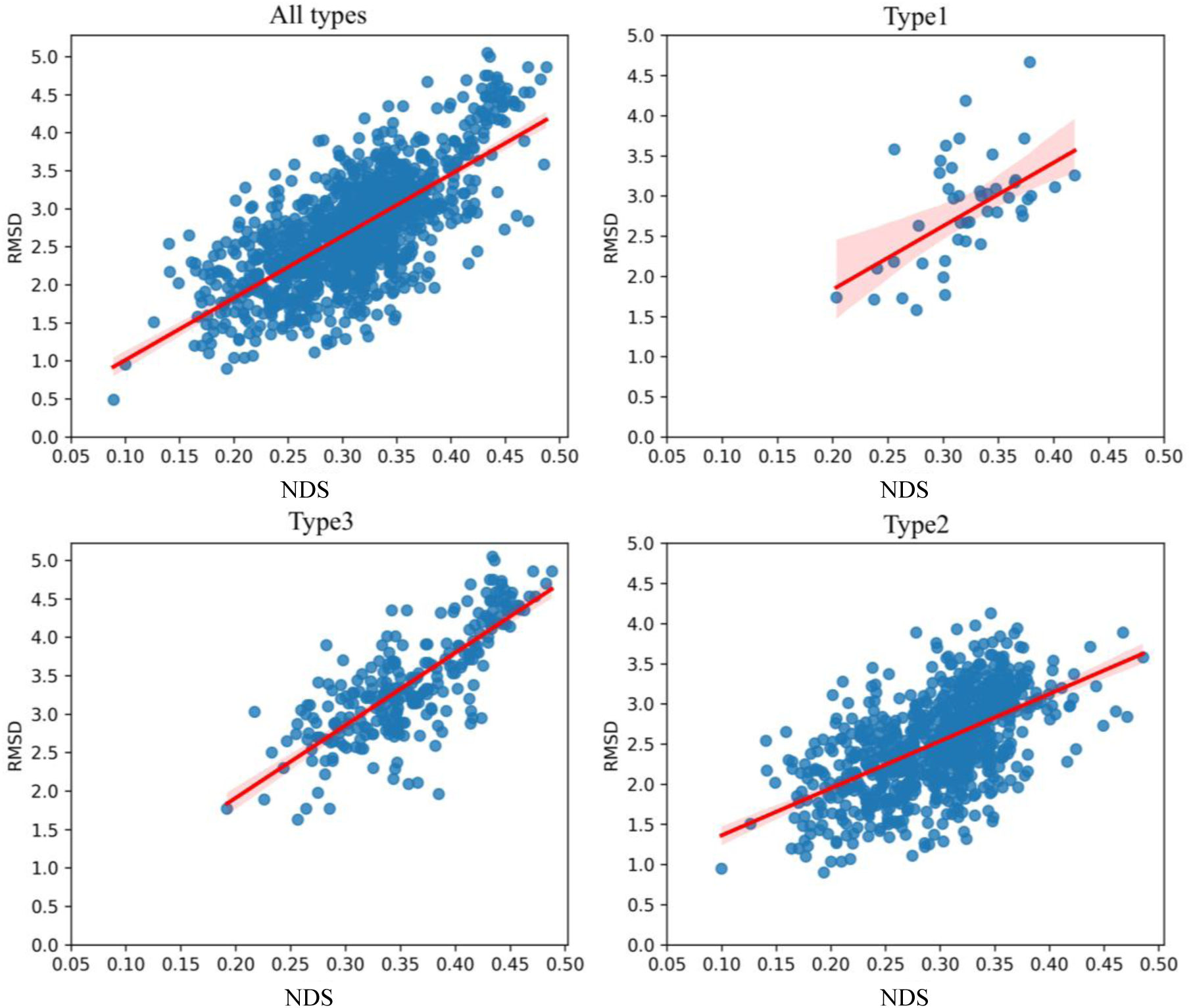
Relationship between network dissimilarity. (NDS) and domain backbone deviations (RMSD). The straight lines are linear regression lines to infer how linear the relationship is.

**Table 3.** Statistics of Network Dissimilarity. Linear correlation between network dissimilarity (NDS) and backbone structural deviation (RMSD).

| Types | Spearman Correlation Coefficient | Linear regression equation |
| --- | --- | --- |
| Type1 | 0.475 | $Y=7.905X+0.314$ |
| Type2 | 0.572 | $Y=5.753X+0.805$ |
| Type3 | 0.741 | $Y=9.213X+0.08$ |

We next compared the components of NDS, that is EDS, EWCS and CRS to better understand the local and global network differences among domain pairs. When we performed principal component analysis to understand the distribution in a reduced dimension and relationship of each component, we observed that EWCS and NDS have a high positive correlation, while other components have no or very weak negative correlation to NDS (Supplementary Figure S10). We also observed significant differences of EWCS and CRS distributions among domain comparison types, but some statistically insignificant differences for EDS (Figure 5B, 5C). Type3 domain pairs also have significantly lower CRS and higher EWCS values, suggesting improper alignment of local node clusters and significant clusters changing of interacting node residues. These observations point to the fact that the network dissimilarities among domains arising from different domain combinations are mostly due to the combined elevated effect of the local node clustering changes, due to the differences in local non-covalent interactions (EWCS) and poor global alignment of these local node clusters (CRS) to some extent.

### 2.4. Case studies

To understand the above patterns in detail, we picked up a few pairs for a detailed study to elucidate the intricate principles behind these trends. We sought to find domain superfamilies which have been paired up in all three types of domain comparisons. Surprisingly, we only found two superfamilies for this criterion, CATH homologous superfamily 3.40.50.2300 (Response regulator) and CATH homologous superfamily 2.60.40.10 (Immunoglobulins).

The superfamily Response regulators (CATH:2.60.40.10) comprise domains with Rossmann fold which has an αβα sandwich topology. This superfamily contains around 2543 domains, which are again distributed in 1676 functional families (FunFams), annotated with 32 unique EC and 1325 unique GO terms in CATH. Several proteins in our dataset contain this superfamily which is evident from Figure 1. We considered the following PDB IDs for further case studies: 1Z17, 3UK0 and 4G97, which contain at least one domain from this superfamily. The structure with PDB ID 1Z17 belongs to a branched amino acid (Leu/Ile/Val) binding protein (otherwise known as LIVBP or LivJ) in the periplasm, which helps to transport nutrients across the membrane (Trakhanov et al., 2005). This protein has two domains from the same superfamily of response regulators and is bound to Isoleucine in its interdomain cleft in this structure. LIVBP holds the nutrient ligand Ile using its dominant N-terminal domain, with the help of comparatively more non-polar interactions, along with a few polar interactions from less dominant C-terminal domain. The structure with PDB ID 3UK0 belongs to a solute binding protein (SBP), a member of ATP-binding cassette (ABC) transporters family and plays a specific role in binding and importing aromatic compounds derived from lignin (a major carbon source for microorganisms) degradation (Tan et al., 2013). Like LIVBP, this protein harbours a pre-bound ligand 3-(4-hydroxly) phenylpyruvic acid (HPP), in the cleft between two domains, where both the domains are again from the same response regulator superfamily (CATH: 3.40.50.2300). The binding site cleft is largely hydrophobic, with additional hydrogen bonding for specificity. The domains of both the proteins (PDB ID 1Z17 and 3UK0) lack higher sequence identity and uniformity in sizes, though very similar in the overall fold. Apart from this, when several SBPs (or PBPs) are classified based on their structure and sequence, it was found that these two proteins are members of the same class of SBPs (Class I) according to Fukami-Lobayashi (Fukami-Kobayashi et al., 1999) scheme and also belong same cluster (Cluster B) according to Berntsson structural classifications (Berntsson et al., 2010; Tan et al., 2013). The third domain structure (PDB ID 4G97) belongs to a protein PhyR, a phosphorylation dependent stress response regulator in bacteria (Herrou et al., 2010). This protein contains two unique domains from two different superfamilies, where the N-terminal domain belongs to PhyR, sigma-like (SL) domain superfamily (CATH: 1.20.140.160). The active site in C-terminal domain (receiver domain), receives phosphorylation signals upon stress which induces conformational changes in the receiver domain, disrupting the interactions with the N-terminal SL domain, allowing to go to an active state and induce subsequent regulations of gene expression. It should be noted that the response regulator superfamily domain in 4G97 interacts with the cognate domain in a different orientation than other two proteins in this case study. The domain pairs forming different types of domain comparisons is shown in Supplementary Figure S11A.

The superfamily Immunoglobulins (CATH:2.60.40.10) comprises domains with immunoglobulin-like fold which are characterized by two-layer beta-sandwich of beta strands arranged in two beta-sheets. This superfamily contains around 31905 domains, which are again distributed in 6111 FunFams, annotated with 77 unique EC and 4604 unique GO terms in CATH. Despite higher numbers of members, our dataset only contained five domains arising from four proteins (Figure 1). We considered the following PDB IDs: 2FBO,1UT9, and 3RX7 to represent this superfamily. The structure with PDB ID 2FBO belongs to variable region-containing Chitin binding protein 3 (VCBP3), which has a total of three domains, two N-terminal V-type immunoglobulin domains (V1 and V2) followed by a C-terminal chitin binding domain (Prada Hernández et al., 2006) out of which first two domains are crystalized in this structure. This protein helps in immunorecognition using V-type immunoglobulin domains and chitin binding using the chitin binding domain. The V1 and V2 domains have very low sequence identity, however their exceptional structural similarity suggests strong selection pressure to maintain their topology for immune recognition of antigens. The PDB ID 1UT9 comprises two domains from a multi-modular glycoside hydrolase Cellobiohydrolase CbhA that plays a role in cellulose degradation (Schubot et al., 2004). Similarly, another cellulase protein (AaCel9A) with PDB ID 3RX7 (Moréra et al., 2011) was used which also has similar modules like 1UT9, a N-terminal Ig module and a C-terminal catalytic domain. The catalytic domain (CATH: 1.50.10.10) is responsible for hydrolytic activity on carbohydrate chains, while the function of Ig domain is still unknown. It is believed that Ig module is involved in structural and functional integrity of the protein using large number of hydrophobic and hydrophilic interactions between two domains, and entire or half loss leads to complete loss of enzymatic activity (Béguin & Alzarit, 1998; Kataeva et al., 2004; Schubot et al., 2004). It is also noteworthy that these two proteins belong to the same family of Glycoside Hydrolases (GH9) in the CAZY database (Drula et al., 2022), where the former has an exonuclease activity while the later has an endonuclease activity. A summary of the combination of domains used for comparison is shown in Supplementary Figure S11B.

With the help these domain pairs, we first investigated the degree variability between homologous domains. We considered a degree difference of five between topological equivalent residues to define higher degree variability, as many such studies use a cut-off much greater than five (Lopes et al., 2021) or close to five (Prabantu et al., 2021). Moreover, this difference is an empirical value to infer higher variability. From Table 4 and Supplementary Table S2, we observed that the higher degree difference is associated with a more remarkable change in relative solvent accessibility. Most of the topologically equivalent residues with lower degree have moved to a solvent accessible surface in the domain, while the equivalent higher degree node remains in buried or moderately buried (if RSA cutoff is 20%) regions of domains. In a few instances, the change in solvent accessibility is accompanied or compensated by change in number of atoms in the residue (at least 40% difference in number of atoms), thereby changing the size of the residue. The variation of degree among topological equivalent residues of homologous domains can be assumed to be due to changes in orientations that move the residues to the solvent exposed regions and the change in size of the residue in that region (Figure 7), which could be associated with the functional and structural significance of the specific domain. The flipping of residues to the surface also could be due to the entropic advantages associated with those residues in the other domain. We also noticed that variation in degrees of residues arise when their corresponding interacting residues could not be mapped onto the homologous domain, and this gap in mapping arises due to poor alignments. Interestingly, we also observed relationships between degree differences and clustering coefficients of the residues. Clustering coefficients is a measure of clustering of nodes in local regions, where a higher value suggests a well knitted neighbourhood of the node. We observed that the residues with lower degree than the equivalent residue in homologous domain have a higher clustering coefficient in many instances. For example, Y49 in N-terminal domain has a degree of 12, while the topologically equivalent residue D173 has a degree of 6. From Figure 8, it is clear that the residue with lower degree, has better knitted interactions with its neighbouring residues than the higher degree residue in remote homologous domain. This observation hints towards the compensations of information flow by well-connected neighbouring residues rather than higher self-interactions, giving a higher degree. A few previous studies (Barabási & Oltvai, 2004; N. Wang et al., 2021) also observed these relationships between clustering coefficients and degree, implying a universality of this observation. When the degree variability is compared across domain comparison types, we observed that type3 domain pairs that arise from proteins of different domain combinations, have plenty of such residues whose solvent accessibility, amino acid size differs, and many interacting residues could not be mapped when compared with corresponding amino acids from the other domain in the pair. Therefore, showing higher degree variability as seen in Figure 3A. Type2 pairs though shows similar or lower number of such residues than type1 making this type as the least degree variable (Table 4 and Supplementary Table S2).

**Figure 7.**
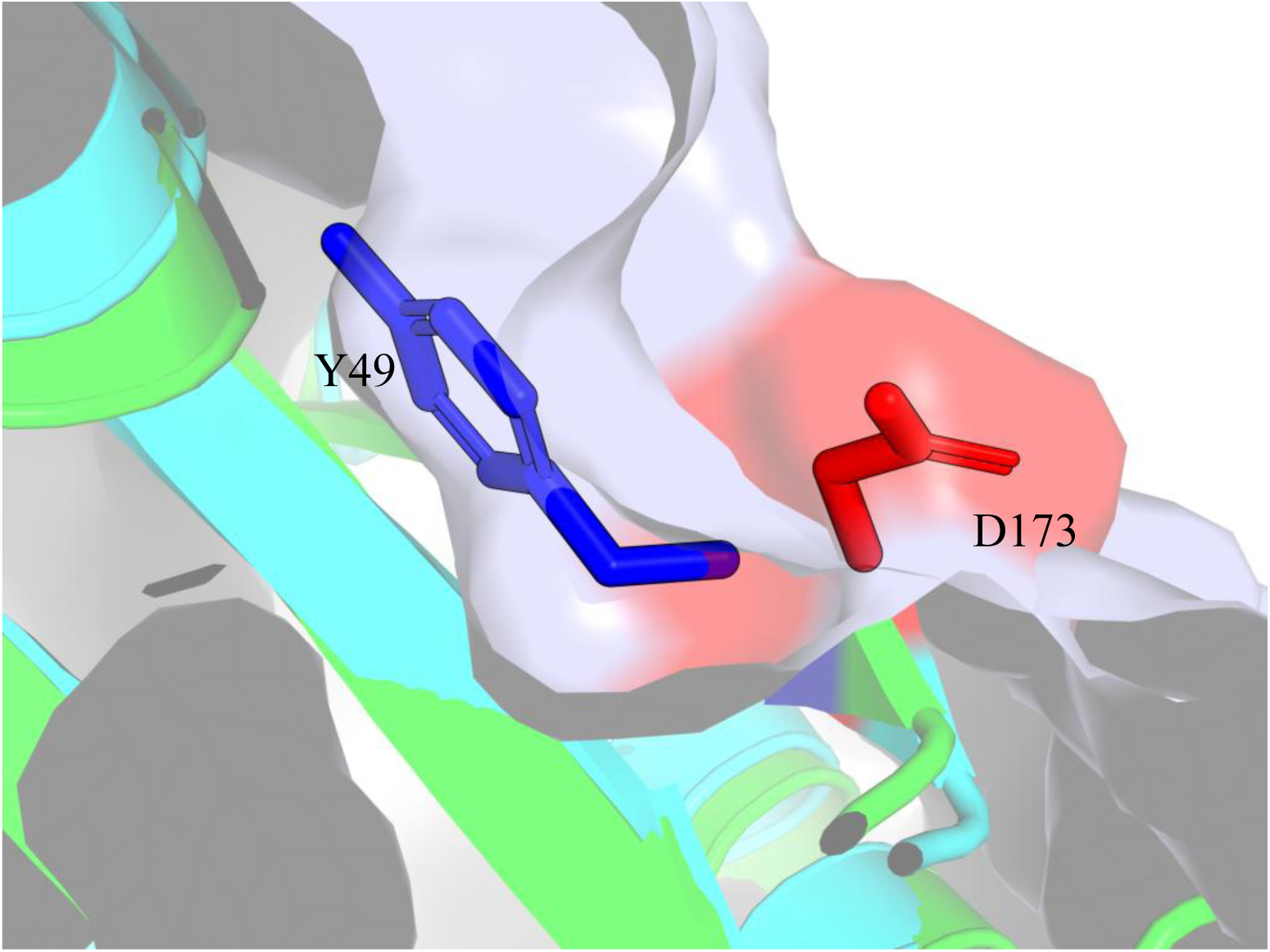
Illustration of solvent accessibilities in remote homologous domains. This snapshot is taken from branched amino acid (Leu/Ile/Val) binding protein (LIVBP), PDB ID: 1Z17. This example is taken from a Type 1 domain comparison from a Response Regulator superfamily (CATH ID: 3.40.50.2300). The blue and red coloured sticks show Tyrosine and Aspartic acid side-chains from N and C-terminal domain of PDB ID 1Z17, respectively. The relative solvent accessibilities are 5.6 and 51.3 Å^2^ for Y49 and D173, respectively.

**Figure 8.**
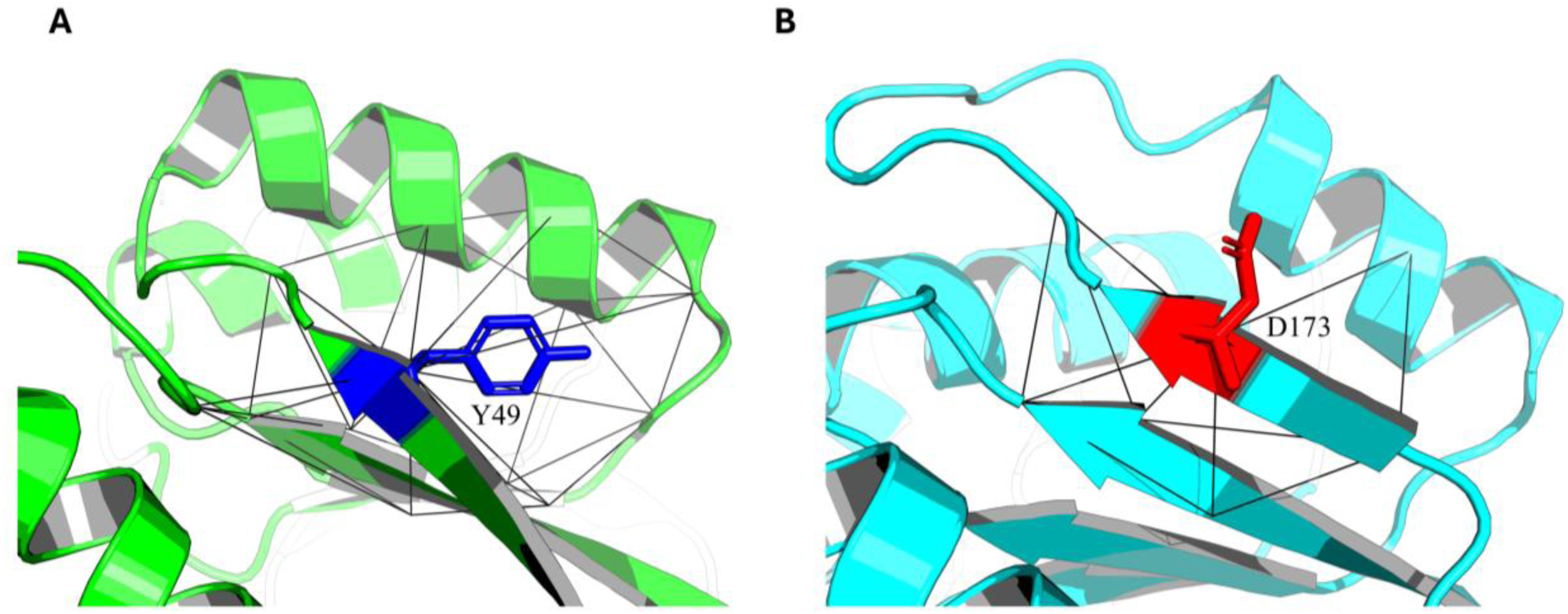
Clustering coefficients of nodes for a Type1 domain pair. Interactions are shown as lines between interacting residues of **A**. Tyrosine 49 of N-terminal domain of PDB ID 1Z17 and **B**. Aspartic acid of C-terminal domain of PDB ID 1Z17. Y49 has 12 interacting residues and makes only 20 edges among interacting partner residues, giving rise to clustering coefficient of 0.3. D173 has six interacting residues and makes six edges among interacting partner residues, giving rise to a higher clustering coefficient, 0.4.

**Table 4.**
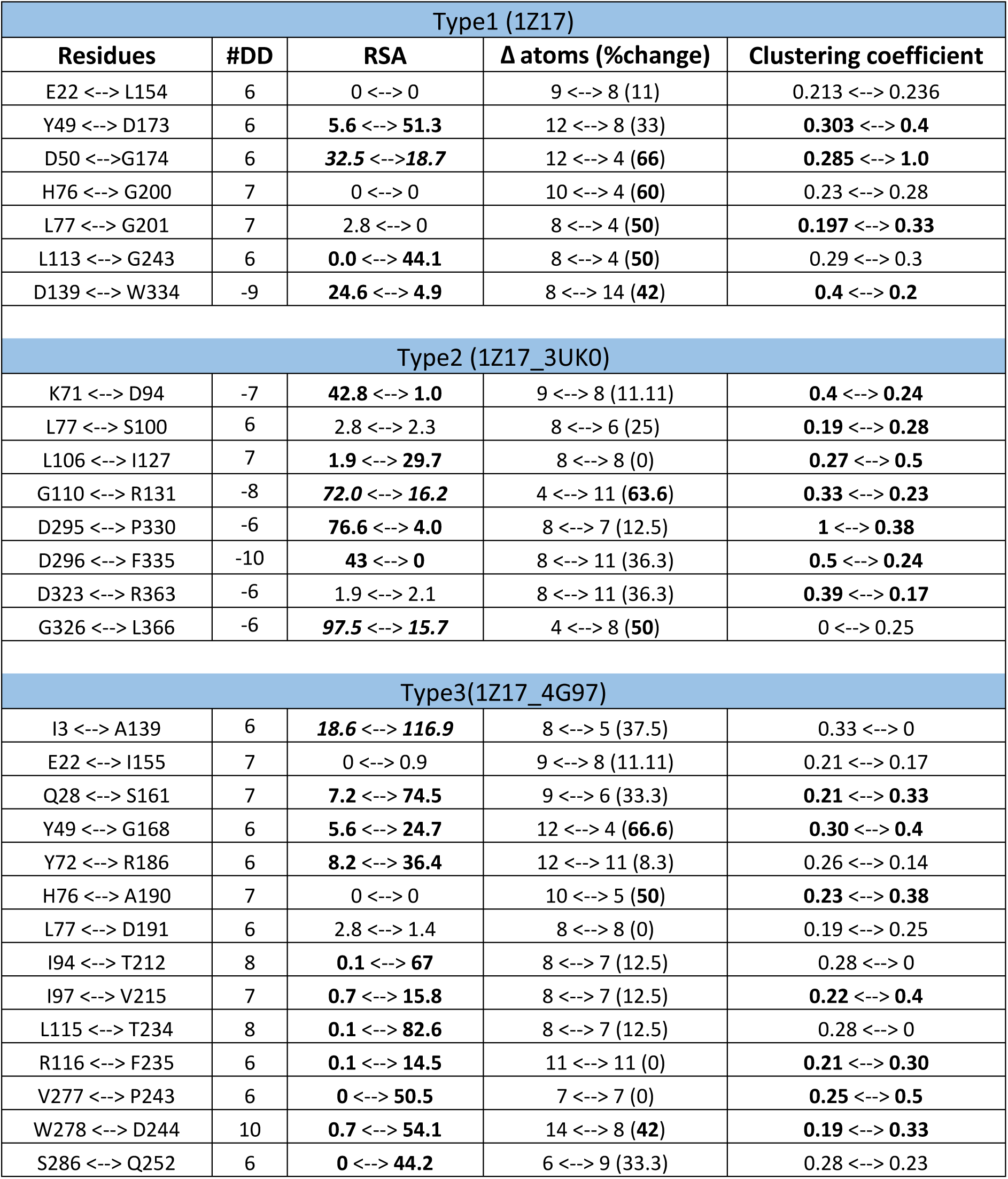
Properties of amino acid residues showing high degree differences in superfamily 3.40.50.2300 (Response regulator). #DD represents the difference of degrees between residue pairs. The degrees are substracted from left to right and a negative sign indicates the right side residue with a higher degree. The properties to look for is shown in bold and satisfies our hypothesis. In some instances it is shown in both bold and italic font when a relaxed solvent accessibility cutoff is used to show buridness. **RSA**: Relative solvent accessibility. The percentage change in number of atoms was calculated by (max(no. of atoms)-lowest(no. of atoms))*100/max(no. of atoms). Wherever the change in atom number exceeds 40%, it is shown in bold. For clustering coefficients, if the difference is around 0.1, it is shown in bold font.

We also looked into the distribution of hubs in each of these domains (Supplementary Table S3). We observed that the number of non-conserved hubs is higher than conserved hubs in all pairs in the case study. For superfamily Response regulator, Type3 domain pairs showed a 1:4.2 ratio of conserved to non-conserved hubs while Type1 and Type2 showed 1:2.6 and 1:1.8 respectively, elaborating the observed pattern of higher rate of change of non-conserved hubs with respect to conserved hubs in Type3 domain pairs (Figure 4). As Type3 pairs showed better correlation coefficient of types of hubs, we fit the number of conserved and non-conserved hubs into the linear regression equation, and we observed that the number of non-conserved hubs can be closely deduced using number of conserved hubs. Although, the rate of change of hubs for Immunoglobulins superfamily follow a similar pattern, Type1 domain pair (2FBO) showed a little elevated rate of change in hubs despite having higher structural similarity as the C-terminal domain looks to have loose interactions and therefore a smaller number of hubs. This makes the number of hubs fall below the linear regression line and hence a small deviation from the probable linear relationship.

Figure 7 and Supplementary Figure S12 shows that the core of the domain structure is maintained by a few hubs (black spheres) thereby maintaining the fold of the homologous domains, while unique hubs (red and blue spheres) provide necessary structural and/or functional support to important regions evolved for its function. This approach demonstrates how the differences can be captures well in structural connectivity between homologous domains. It is also observed that one of the domains in Type3 pairs to have fewer unique hubs than the other (26 vs 4 unique hubs in Response regulator superfamily). This also could be due to the flexibility (lower degree) of peripheral regions of domains when compared with domains from different domain combination (Type3). This is possibly due to the different extents of evolutionary pressures on domains in multi-domain proteins for easy modification (mutation of non-hubs) of residues to adapt new function. Despite having different domain combinations and protein function, the binding/active site the domains belonging to same homologous superfamilies is located in the same region in the tertiary structure (Figure 7). The unique hubs are located accordingly to harbour a specific ligand and also based on the dominant involvement of the domain. For Type3 domain pairs, the hubs are only located in the core region when both domains are aligned and mostly no hubs around the peripheral regions. This clearly indicates the poor alignment of the peripheral regions leads to lower topological equivalent position and hence less comparable hub detection. We think that these additional structural embellishments have co-evolved along with the cognate interacting domain for an evolved function with a different quantity of evolutionary pressure. When investigated for secondary structure propensity of these hubs, we did not observe any patterns. In other words, these hubs occur in major secondary structures as well as in loops connecting them. However, conserved hubs showed minor preference to be present in secondary structures (like strands and helices) as observed in the above case studies.

**Figure 7.**
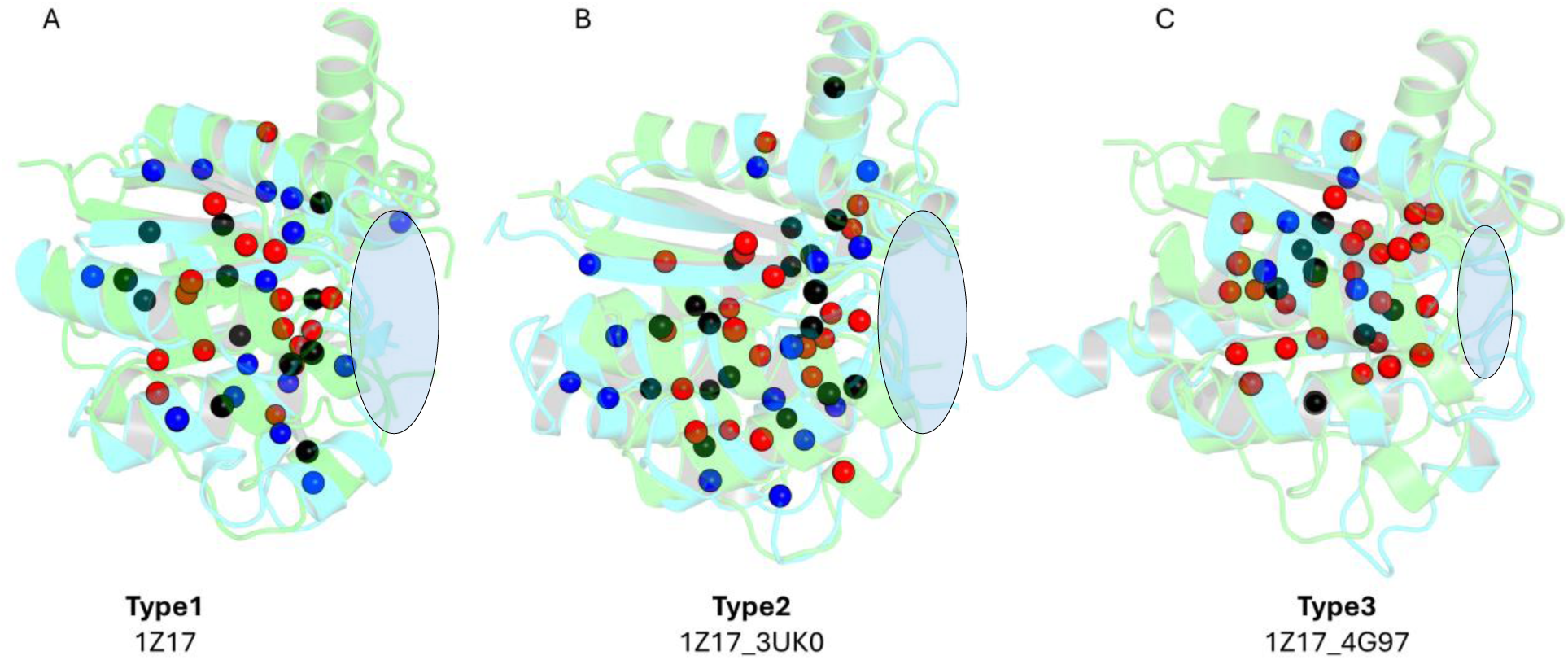
Distribution of hubs in superfamily Response regulator (CATH: 3.40.50.2300). Homologous superfamily domains in types of comparison, shown in A, B, C are superimposed where one domain is coloured green and the other cyan. Black spheres show conserved hubs between the two networks. Red spheres show unique hubs to domain1 (green cartoon) and blue spheres show unique hubs to domain2 (cyan cartoon). The oval shapes show the binding/active site in the domain.

**Figure 8.**
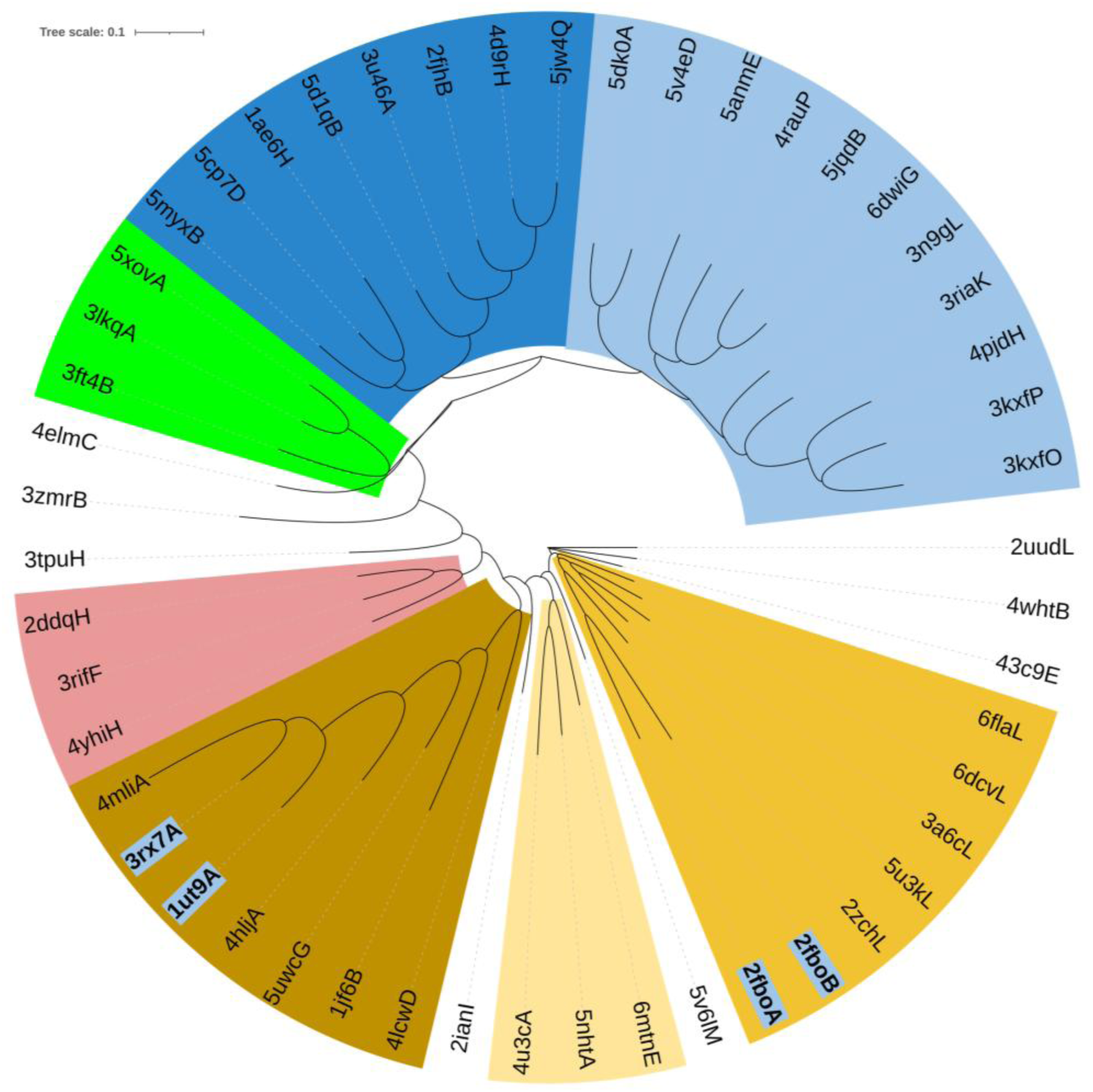
Phylogenetic dendrogram of Network dissimilarities of homologous superfamily domains from Immunoglobulins. The case studies are shown in bold font and light blue background. Clades are coloured based on visual clade formation.

From all these observations, we obtained some information about the evolution of these domains. So, we explored their phylogenetic relationship. Firstly, we used ECOD database (Cheng et al., 2014) which contains domains of evolutionary relationships. We found that the domains in Type1 and Type2 comparisons belong to same T-groups, implying homologous domains with similar topological connections which have evolved in the same rate. On the other hand, Type3 domain pairs have some level of structural changes due to evolution which groups them in different T-group, implying homologous domains but distinct topologies. Secondly, we used the network dissimilarity scores (NDS) of several single domains of same homologous superfamily extracted from multi-domain proteins (taken from CATH database again and not present in the current dataset) to compare the phylogenetic relationship of networks wiring the domain structure. From Figure 8 and Supplementary Figure 13, we observed that domains in Type1 and Type2 are in a clade, and they share similar network divergence. Type3 domains are present in different clades and distance between them shows larger network divergence in course of evolution. Interestingly, although Type1 comparison contain two domains of superfamily Response regulator in a single chain, both domains share very early evolutionary ancestor as one of the domains have additional secondary structures, leading to additional network components. Apart from this peculiarity, the phylogenetic study shows that when domains of same domain combinations proteins are compared, their network divergence is minimal and share a very recent ancestor, while domains from two different domain combinations have diverged a lot to adopt a completely different function.

Additionally, we also investigated the similarities of functional sites and a few network parameters between homologous domains. We observed that most of the functional sites of the in each type of domain comparison align well with another functionally relevant and topological equivalent site in the other domain (Supplementary Table S4). Wherever the sites are not reciprocated, it was observed that the corresponding nodes have very minimal degree differences. A significant deviation is observed for Type3, which are mostly due to poor mapping of interacting residues of each node, as mentioned earlier. This suggests that most of the functional sites are conserved among homologous domains. When these functional residues were mapped onto domain structures in Figure 9, it was observed that these residues in Type 2 domain comparisons align well and maintain the properties needed for a similar function of the protein. On the other hand, in Type 3 domain comparisons, these functional active site residues are re-oriented and re-positioned. One such example is shown in Figure 9, where a residue (F276) important for the hydrophobic pocket in one of the domains is re-oriented (N242) in the other domain. This re-orientation in the domain (2nd domain of 4G97) is rather used in the phosphoryl transfer pathway in the protein instead of maintaining the hydrophobic core as seen in the other domain (1st domain of 1Z17). These fine-tuned structural changes are mostly located in the loop regions of the domain, which evolve faster than strands and helices (Panchenko & Madej, 2005; Schaefer et al., 2010; Siltberg-Liberles et al., 2011), so as to accommodate a newer function in Type 3 domain comparison, instead of accommodating a similar function in Type 2 domain comparison.

**Figure 9.**
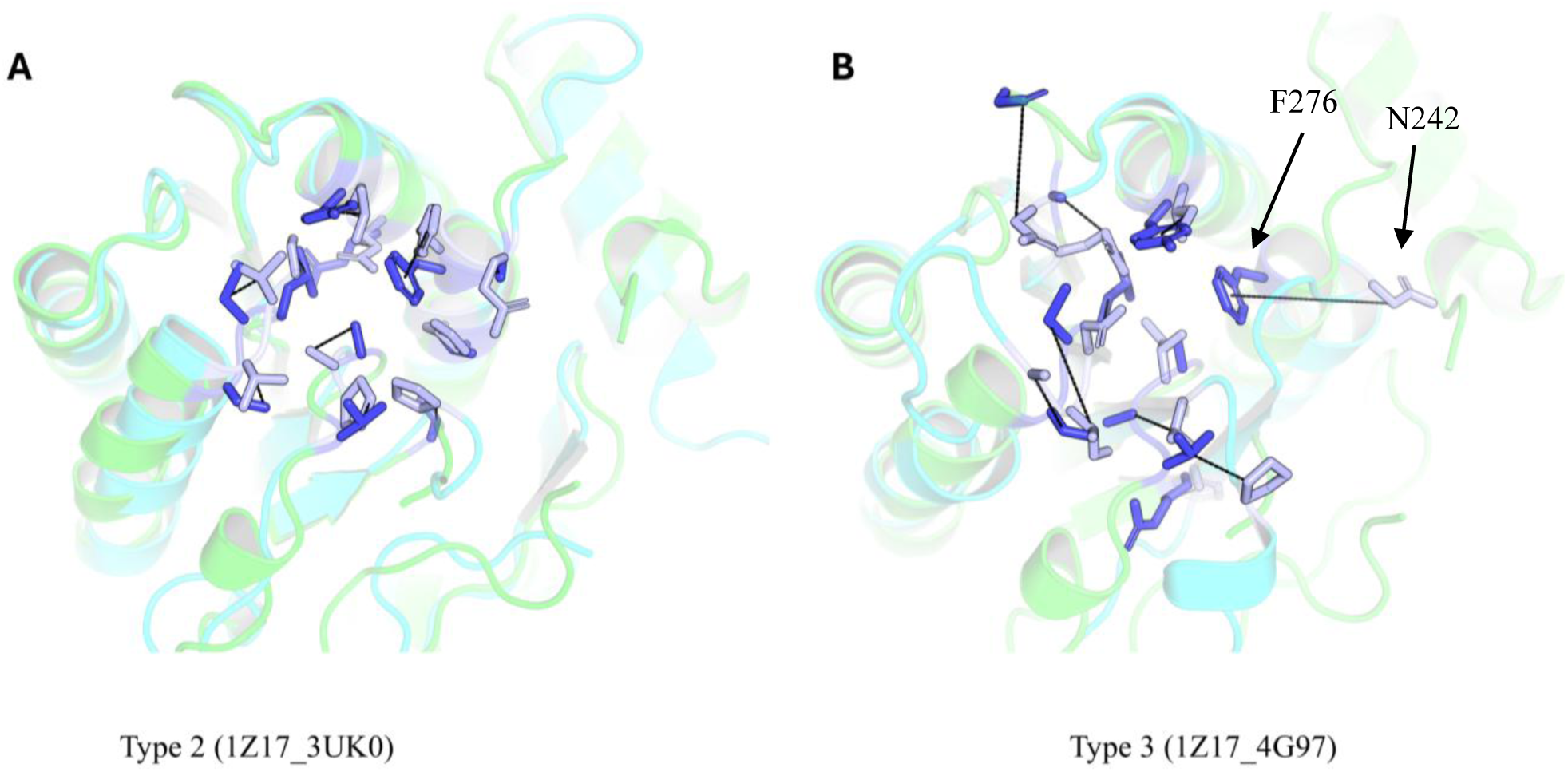
Mapping of functional residues on structures. The functional residues of each domain in each type of domain comparison are mapped. Green and cyan colour cartoons represent 1^st^ and 2^nd^ domain in the domain comparison. Here, green colour cartoon shows 1^st^ domain of PDB ID 1Z17. The functional residues and their topological equivalent residues in the other domain are shown in stick representation. Blue sticks represent functional residues or topological equivalent residues in the 1^st^ domain of PDB ID 1Z17. Light blue sticks represent functional or topological equivalent residues in the 1^st^ domain of PDB ID 3UK0 (A) and 2^nd^ domain of PDB ID 4G97 (B). The dashed lines show the difference in position and orientation of these residues in two domains. Single headed arrows point to the residues which are reoriented.

Several unique hubs are distributed over the structure mostly to provide structural or functional support for the tertiary structure while their topological equivalent residues in the other domain are functionally or structurally not important. Moreover, when we compared a few more network parameters for the above case studies, we observed unnoticeable differences between all types of domain comparisons. In other words, we could not establish any pattern (Supplementary Table S3) for those network properties except mentioned here, as parameters do not show any particular bias for any type of domain comparison as seen in these case studies.

## 3. Conclusions

The homology of domains is generally inferred from the sequence identities. But it is also observed that there are several domains with diverse sequences while maintaining a very similar structure, and sometimes similar functions. Such domains are grouped in a homologous superfamily in domain databases like CATH, which implies a very common ancestor among the members of the group. It is also observed that the relationship between evolutionary relationship, the structural fold and the functions they do are not straightforward. Moreover, multidomain proteins with similar domain architecture are very related. In this study we incorporated network biology using interatomic non-covalent interactions to compare remote homologous domain and the influence of domain interactions arising from another domain of different structural origin. In other words, we aimed to investigate the intricate structural differences lining highly similar structural domains when shared domain from different domain architectures are compared.

We created a dataset of two domain proteins to reciprocate multi-domain behaviour and segregated them into different types of domain comparisons. We found that our dataset follows many universal laws of domain grammar, like power-law, divergent sequence-structure relations, limited combinations of domains etc. We compared the networks of these domains in pairs in both local and global level. At local level, we compared the degrees of topological equivalent residue among domain architectures and found that remote homologues from different domain architectures tend to have higher degree variabilities implying local interaction variations. These local variations in structure also alters the global backbone of the domain to some extent. We also found that all remote homologous domains adopt a different approach for their structural integrity and this variation in structural integrity again varies based on the domain architecture under comparison. The structural cores of such domains are maintained by a few important residues, while other highly interacting residues help each domain for structural embellishment to adopt a newer function. In global network level, we compared the network dissimilarity scores (NDS) which are shown to be a robust scheme for structural comparisons. We noticed that network dissimilarities are mapped in a hierarchical manner by nature depending on the relationship of the pairs in comparison. Multiple conformations of single domain proteins and homologous single domain proteins take the lower positions in this hierarchy, while remote homologous take the higher positions. Furthermore, network variations are found in remote homologous when similar or different domain-domain interactions are found in terms of domain combinations. Domains within a multi-domain architecture and similar domain architectures reorient the local interactions to minimize global structural differences. These network differences arise due to fine-tuned re orientations and re-positioning of amino acids in the remote homologous domain, which are mostly essential for adaptation of a new function. These fine differences would not be apparent through sequence alignment measures or RMSD values.

Additionally, we took some case studies to better understand these patterns. We found that the change in solvent accessibility, change in size of the residues and improper mapping of interactions onto the partner domain in the comparison, are the major reasons for degree variability. Remote homologues from different domain combinations show these properties extensively, mostly due to poor alignments. Interestingly, we also found that reduction in degree of residues is compensated by higher clustering coefficients, suggesting compensations of information flow by a well-connected neighbourhood. These results points to a question that how evolution has impacted these observations. We used network evolution of multiple domains of the same superfamily to find how evolution impacts the trends observed in this study. We found that, structural networks of remote homologues within a domain architecture or from a same architecture are less diverged and share a very recent ancestor. On the other hand, remote homologues from different domain architectures are highly divergent in their structural network and share a very early ancestor. This clearly suggests that there is a significantly different evolutionary pressure on domains in multi-domain proteins of different domain combinations and these domains are in the verge of entering or exiting a new evolutionary curve for new functions.

These results strongly imply to shift the definition of homology among domains from only sequence, structure and functional similarity towards involving network similarities. Apart from this, our work provides a report of structural comparisons of remote homologues and the effect of domain interactions arising from an interacting domain in multi-domain architecture. We strongly believe that this work will pave way for more accurate functional annotation transfer.

## 4. Materials and methods

### 4.1. Dataset creation

We have employed interface library (Sen & Madhusudhan, 2022) for creation of the dataset. This library contains protein chain-chain and intrachain domain-domain interfaces. We only used domain-domain interfaces (around 66442 interfaces) from this library to represent interacting domains. RCSB PDB (Berman et al., 2000a) was mined to filter monomeric proteins with good structural information using filters (such as, protein as polymer type, only one polymer instance and number of protein chain instance per assemble, monomer in its oligomeric state, X-ray crystallographic experimental method, refinement resolution of better than 3Å with R_free_ and R_work_ of 0.3 and 0.25 respectively. Despite these filters, in some PDB IDs, proteins exist both as monomers and multimers in their biological assembly (for example, Fructose-1,6-bisphosphatase 1 with PDB ID 1NUW which exists as both monomer and homo-4-mer in its biological assembly), and such entries were removed. A few protein structures which were not annotated to UniProt (Bateman et al., 2023) (for example, iron uptake ABC transporter binding protein with PDB ID 4JCC), would give ambiguity while studying them in depth, were removed from the study. Finally, 5072 monomeric protein PDB IDs were used against PISCES server (G. Wang & Dunbrack, 2003) to recognise redundant proteins by using further filters (a maximum sequence identity of 40% among them, maximum resolution of 3Å, and other default parameters) leading to 922 non-redundant monomeric proteins. We consulted CATH (Orengo et al., 1997) database for structural domain definitions, which is in line with the interface library used here. The domains with synthetic constructs, whose boundaries start with a negative residue number, were re-formatted to start from residue number one and we considered only two domain proteins for this study. pdb-tools (Rodrigues et al., 2018) was employed to perform necessary structural editing of proteins into domains. We used NOXclass (Zhu et al., 2006) to predict domain interactions to permanent and transient interactions at a 70% cutoff, resulting in total of 375 monomeric two domain proteins. Later, we employed a mutation filter using RCSB to remove proteins with engineered mutations (please see later). The resulting 294 proteins in the dataset were used in the study.

### 4.2. Sequence and structure alignments

We extracted the sequences of domains from the PDB file using pdb_tofasta module in pdb-tools (Rodrigues et al., 2018). The sequence identities were realised by aligning them using MUSCLE (version 3.8.31) (Edgar, 2004) and using the alignment in ClustalW2 (version 2.1) (Larkin et al., 2007).

Structural alignments of domains were done using TM-align (Zhang & Skolnick, 2005) and a length normalised TM-score was used to infer the quality of superposition. The aligned positions were considered as topological equivalent positions of amino acids in domains.

To have meaningful comparisons between domains, we employed a TM-score cutoff of more than 0.5 and structural alignment length more than 50% of the size of the smaller domain.

### 4.3. Protein structural network construction and comparison

Protein structural network (PSN) is a representation of protein or domain in the form of nodes and edges, that preserves both topology and the non-covalent interactions within them. Here, we denoted the amino acid residues as nodes of the network and the relation between the nodes as edges to make an undirected network. These edges can be any relationship between the nodes, and here, the relationship is the non-covalent interaction between atoms of the residues. We used the number of atomic interactions (both backbone and sidechain atoms) arising from spatial proximity to weight the edges formed by the residues. In other words, the edge weights (I_ij_) are defined as the ratio of actual number of atomic contacts between two sequentially non-adjacent residues (n_ij_) to the maximum number of atomic contacts between such residue types within the whole dataset (N_ij_) (Equation 1). The spatial proximity of atoms was inferred by a distance cutoff of 4.5Å (related to the sum of atomic radii (Heringa & Argos, 1991)) which captures explicit atomic connectivity. The maximum possible atomic contacts between all residue types are given in Supplementary Figure S2. These edge weights range from zero to one for weak or no interaction to very strong interactions, respectively. The edge weighted undirected network is stored as an adjacency matrix (A) for further comparisons.

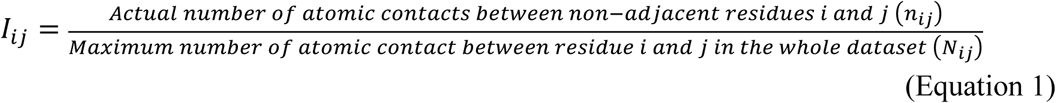

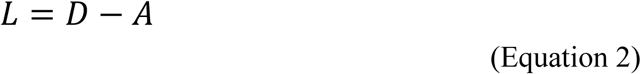

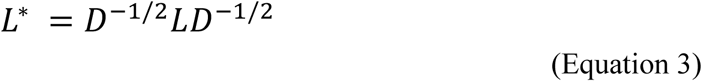

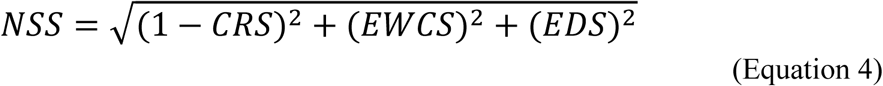

An in-house python script was used to compare the domain networks using spectral decomposition. Topological equivalent residues were only used for network comparisons to compare networks of equal size. The degree of a node in a network denotes the total number of edges that node has. The above adjacency matrix (A) was converted to Laplacian matrix (L) using equation 2, and resulting Laplacian matrix was normalised using diagonal degree matrix (D) given in equation 3. The normalized Laplacian matrix (L*) was then decomposed into its component eigen vectors and eigen values, and network comparisons were carried out by aligning the vectors. A network similarity score (NSS) was obtained by combining different components (Equation 4) which portray both local and global similarities such as edge difference score (EDS), eigenvalue weighted cosine score (EWCS), and correspondence score (CRS). More details on the spectral decompositions and significance of its components are explained somewhere else (Gadiyaram et al., 2017, 2019). The NSS spans a value of 0 to 1.73, where a value of zero implies identical residue networks. Therefore, the term Network Dissimilarity Score (NDS) is used instead of NSS, where a higher value would mean dissimilarity between networks.

### 4.4. Phylogenetic analysis

We considered several CATH domains belonging to the superfamilies for case studies. To reduce computational burden, only 50 domains per case study were considered, which also includes the domains in the case studies. Pairwise network similarity/dissimilarity scores in a distance matrix form was employed for inferring evolution among the domains. The matrix was first converted to phylip (Cummings, 2004) format to be used in FastME 2.0 (Lefort et al., 2015). Algorithm TaxAdd_BME (Desper & Gascuel, 2004) and tree refinement with SPR (Hordijk & Gascuel, 2005) was used for drawing trees. In addition, a Newick format was exported to be used in other visualization tools like iTOL (Letunic & Bork, 2024). The clades were coloured based on visual interpretation from the dendrogram to represent similar residue networks.

### 4.5. Solvent accessibility and network parameters

Solvent accessibility of residues was calculated using NACESS (S.J. Hubbard & J.M. Thornton, 1993). The relative solvent accessibility (including all-atoms) was used to infer the same. A cutoff of 7% RSA was used to define completely buried residues. In some instances, a relaxed cutoff of 20% was used to define partially buried residues.

Network parameters were calculated using NetworkX (Hagberg et al., 2008) package in python.

All the structural Figures were generated using Pymol (The PyMOL Molecular Graphics System, Schrödinger, LLC).

### 4.6. Principal component analysis

NDS values, including its component scores, such as CRS, EWCS, and EDS, were used to reduce dimensionality into two principal components (PC1 and PC2). A python script was used to perform the analysis and plot the results.

## Acknowledgments

The authors thank lab members for helpful discussions. SSPD thanks CSIR and Indian Institute of Science for financial support. SSPD also thanks Molecular Biophysics Unit of Indian Institute of Science, Bangalore for infrastructural facilities. RS acknowledges funding and support provided by JC Bose Fellowship (JBR/2021/000006) from Science and Engineering Research Board, India and Bioinformatics Centre Grant funded by Department of Biotechnology, India (BT/PR40187/BTIS/137/9/2021). RS would also like to thank Institute of Bioinformatics and Applied Biotechnology for the funding through her Mazumdar-Shaw Chair in Computational Biology (IBAB/MSCB/182/2022).

## Author contributions

NS and RS conceived the idea and concept for the work. SSPD, VMP and RS designed the work. SSPD carried out literature review, data curation, all analyses and writing the manuscript. RS shaped and finalized the manuscript.

## Data availability

The data that supports the findings of this study are available in the supplementary material of this article. The codes and dataset is available in a public repository and can be accessed at https://github.com/Sidhanta-SPD/Domain-RIN-effects-of-domain-architectures

## Abbreviations

PSN: Protein Structural Network
RMSD: Root Mean Square Deviation
NSS: Network Similarity Score
NDS: Network Dissimilarity Score
CRS: Correspondence Score
EWCS: Eigenvalue Weighted Cosine Score
EDS: Edge Difference Score
RSA: Relative Solvent Accessibility
GO: Gene Ontology
EC: Enzyme Commission
LIVP: branched amino acid (Leu/Ile/Val) binding protein
SBP: Solute Binding Protein
ABC: ATP-binding Cassette
VCBP3: Variable region-containing Chitin binding protein 3

## References

1. Albert, R., & Barabási, A. L. (2002). Statistical mechanics of complex networks. Reviews of Modern Physics, 74(1), 47. 10.1103/RevModPhys.74.47

2. Apic, G., Huber, W., & Teichmann, S. A. (2003). Multi-domain protein families and domain pairs: Comparison with known structures and a random model of domain recombination. Journal of Structural and Functional Genomics, 4(2–3), 67–78. 10.1023/A:1026113408773/METRICS

3. Barabási, A. L., & Oltvai, Z. N. (2004). Network biology: understanding the cell’s functional organization. Nature Reviews Genetics 2004 5:2, 5(2), 101–113. 10.1038/nrg1272

4. Bashton, M., & Chothia, C. (2002). The geometry of domain combination in proteins. Journal of Molecular Biology, 315(4), 927–939. 10.1006/JMBI.2001.5288

5. Bashton, M., & Chothia, C. (2007). The generation of new protein functions by the combination of domains. Structure (London, England : 1993), 15(1), 85–99. 10.1016/J.STR.2006.11.009

6. Basu, M. K., Carmel, L., Rogozin, I. B., & Koonin, E. V. (2008). Evolution of protein domain promiscuity in eukaryotes. Genome Research, 18(3), 449–461. 10.1101/GR.6943508

7. Basu, M. K., Poliakov, E., & Rogozin, I. B. (2009). Domain mobility in proteins: functional and evolutionary implications. Briefings in Bioinformatics, 10(3), 205–216. 10.1093/BIB/BBN057

8. Bateman, A., Martin, M. J., Orchard, S., Magrane, M., Ahmad, S., Alpi, E., Bowler-Barnett, E. H., Britto, R., Bye-A-Jee, H., Cukura, A., Denny, P., Dogan, T., Ebenezer, T. G., Fan, J., Garmiri, P., da Costa Gonzales, L. J., Hatton-Ellis, E., Hussein, A., Ignatchenko, A., … Zhang, J. (2023). UniProt: the Universal Protein Knowledgebase in 2023. Nucleic Acids Research, 51(D1), D523–D531. 10.1093/NAR/GKAC1052

9. Béguin, P., & Alzarit, P. M. (1998). The cellulosome of Clostridium thermocellum. Biochemical Society Transactions, 26(2), 178–184. 10.1042/BST0260178

10. Berman, H. M., Westbrook, J., Feng, Z., Gilliland, G., Bhat, T. N., Weissig, H., Shindyalov, I. N., & Bourne, P. E. (2000a). The Protein Data Bank. Nucleic Acids Research, 28(1), 235–242. 10.1093/NAR/28.1.235

11. Berman, H. M., Westbrook, J., Feng, Z., Gilliland, G., Bhat, T. N., Weissig, H., Shindyalov, I. N., & Bourne, P. E. (2000b). The Protein Data Bank. In Nucleic Acids Research (Vol. 28, Issue 1, pp. 235–242). Oxford University Press. 10.1093/nar/28.1.235

12. Berntsson, R. P. A., Smits, S. H. J., Schmitt, L., Slotboom, D. J., & Poolman, B. (2010). A structural classification of substrate-binding proteins. FEBS Letters, 584(12), 2606–2617. 10.1016/J.FEBSLET.2010.04.043

13. Björklund, Å. K., Ekman, D., & Elofsson, A. (2006). Expansion of protein domain repeats. PLoS Computational Biology, 2(8), 0959–0970. 10.1371/JOURNAL.PCBI.0020114

14. Bonello, J., & Orengo, C. (2024). FunPredCATH: An ensemble method for predicting protein function using CATH. Biochimica et Biophysica Acta. Proteins and Proteomics, 1872(2). 10.1016/J.BBAPAP.2023.140985

15. Chakrabarti, S., & Sowdhamini, R. (2004a). Regions of minimal structural variation among members of protein domain superfamilies: application to remote homology detection and modelling using distant relationships. FEBS Letters, 569(1–3), 31–36. 10.1016/J.FEBSLET.2004.05.028

16. Chakrabarti, S., & Sowdhamini, R. (2004b). Regions of minimal structural variation among members of protein domain superfamilies: application to remote homology detection and modelling using distant relationships. FEBS Letters, 569(1–3), 31–36. 10.1016/J.FEBSLET.2004.05.028

17. Cheng, H., Schaeffer, R. D., Liao, Y., Kinch, L. N., Pei, J., Shi, S., Kim, B. H., & Grishin, N. V. (2014). ECOD: An Evolutionary Classification of Protein Domains. PLOS Computational Biology, 10(12), e1003926. 10.1371/JOURNAL.PCBI.1003926

18. Chothia, C. (1992). Proteins. One thousand families for the molecular biologist. Nature, 357(6379), 543–544. 10.1038/357543A0

19. Chothia, C., Gough, J., Vogel, C., & Teichmann, S. A. (2003). Evolution of the protein repertoire. *Science (New York*, N.Y.), 300(5626), 1701–1703. 10.1126/SCIENCE.1085371

20. Cohen, E. A., & Barabási, A.-L. (2002). Linked: The New Science of Networks. Foreign Affairs, 81(5), 204. 10.2307/20033300

21. Cuff, A. I., Sillitoe, I., Lewis, T., Redfern, O. C., Garratt, R., Thornton, J., & Orengo, C. A. (2008). The CATH classification revisited—architectures reviewed and new ways to characterize structural divergence in superfamilies. Nucleic Acids Research, 37(Database issue), D310. 10.1093/NAR/GKN877

22. Cummings, M. P. (2004). PHYLIP (PHYLogeny Inference Package). Dictionary of Bioinformatics and Computational Biology. 10.1002/9780471650126.DOB0534.PUB2

23. Dessailly, B. H., Redfern, O. C., Cuff, A. L., & Orengo, C. A. (2010). Detailed Analysis of Function Divergence in a Large and Diverse Domain Superfamily: Toward a Refined Protocol of Function Classification. Structure, 18(11), 1522–1535. 10.1016/J.STR.2010.08.017

24. Drula, E., Garron, M. L., Dogan, S., Lombard, V., Henrissat, B., & Terrapon, N. (2022). The carbohydrate-active enzyme database: functions and literature. Nucleic Acids Research, 50(D1), D571–D577. 10.1093/NAR/GKAB1045

25. Edgar, R. C. (2004). MUSCLE: multiple sequence alignment with high accuracy and high throughput. Nucleic Acids Research, 32(5), 1792–1797. 10.1093/NAR/GKH340

26. Fukami-Kobayashi, K., Tateno, Y., & Nishikawa, K. (1999). Domain dislocation: a change of core structure in periplasmic binding proteins in their evolutionary history. Journal of Molecular Biology, 286(1), 279–290. 10.1006/JMBI.1998.2454

27. Gadiyaram, V., Ghosh, S., & Vishveshwara, S. (2017). A graph spectral-based scoring scheme for network comparison. Journal of Complex Networks, 5(2), 219–244. 10.1093/COMNET/CNW016

28. Gadiyaram, V., Vishveshwara, S., & Vishveshwara, S. (2019). From Quantum Chemistry to Networks in Biology: A Graph Spectral Approach to Protein Structure Analyses. Journal of Chemical Information and Modeling, 59(5), 1715–1727. 10.1021/ACS.JCIM.9B00002/ASSET/IMAGES/LARGE/CI-2019-00002M_0007.JPEG

29. Gerstein, M. (1998). How representative are the known structures of the proteins in a complete genome? A comprehensive structural census. Folding and Design, 3(6), 497–512. 10.1016/S1359-0278(98)00066-2/ASSET/69BEFA4C-E50E-4809-9D97-9799B98D4E82/MAIN.ASSETS/GR4.JPG

30. Hagberg, A., Swart, P. J., & Schult, D. A. (2008). Exploring network structure, dynamics, and function using NetworkX.

31. Halder, A., Anto, A., Subramanyan, V., Bhattacharyya, M., Vishveshwara, S., & Vishveshwara, S. (2020). Surveying the Side-Chain Network Approach to Protein Structure and Dynamics: The SARS-CoV-2 Spike Protein as an Illustrative Case. Frontiers in Molecular Biosciences, 7, 596945. 10.3389/FMOLB.2020.596945/BIBTEX

32. Hegyi, H., & Gerstein, M. (2001). Annotation Transfer for Genomics: Measuring Functional Divergence in Multi-Domain Proteins. Genome Research, 11(10), 1632. 10.1101/GR.183801

33. Heringa, J., & Argos, P. (1991). Side-chain clusters in protein structures and their role in protein folding. Journal of Molecular Biology, 220(1), 151–171. 10.1016/0022-2836(91)90388-M

34. Herrou, J., Foreman, R., Fiebig, A., & Crosson, S. (2010). A structural model of anti-anti-σ inhibition by a two-component receiver domain: the PhyR stress response regulator. Molecular Microbiology, 78(2), 290–304. 10.1111/J.1365-2958.2010.07323.X

35. Hordijk, W., & Gascuel, O. (2005). Improving the efficiency of SPR moves in phylogenetic tree search methods based on maximum likelihood. Bioinformatics, 21(24), 4338–4347. 10.1093/BIOINFORMATICS/BTI713

36. Illergård, K., Ardell, D. H., & Elofsson, A. (2009). Structure is three to ten times more conserved than sequence—A study of structural response in protein cores. *Proteins: Structure*, Function, and Bioinformatics, 77(3), 499–508. 10.1002/PROT.22458

37. Itoh, M., Nacher, J. C., Kuma, K. ichi, Goto, S., & Kanehisa, M. (2007). Evolutionary history and functional implications of protein domains and their combinations in eukaryotes. Genome Biology, 8(6). 10.1186/GB-2007-8-6-R121

38. Iyer, M. S., Joshi, A. G., & Sowdhamini, R. (2018). Genome-wide survey of remote homologues for protein domain superfamilies of known structure reveals unequal distribution across structural classes. Molecular Omics, 14(4), 266–280. 10.1039/C8MO00008E

39. Jin, J., Xie, X., Chen, C., Park, J. G., Stark, C., James, D. A., Olhovsky, M., Linding, R., Mao, Y., & Pawson, T. (2009). Eukaryotic protein domains as functional units of cellular evolution. Science Signaling, 2(98). 10.1126/SCISIGNAL.2000546

40. Kajava, A. V. (2012). Tandem repeats in proteins: From sequence to structure. Journal of Structural Biology, 179(3), 279–288. 10.1016/J.JSB.2011.08.009

41. Kataeva, I. A., Uversky, V. N., Brewer, J. M., Schubot, F., Rose, J. P., Wang, B. C., & Ljungdahl, L. G. (2004). Interactions between immunoglobulin-like and catalytic modules in Clostridium thermocellum cellulosomal cellobiohydrolase CbhA. Protein Engineering, Design and Selection, 17(11), 759–769. 10.1093/PROTEIN/GZH094

42. Keskin, O., Jernigan, R. L., & Bahar, I. (2000). Proteins with Similar Architecture Exhibit Similar Large-Scale Dynamic Behavior. Biophysical Journal, 78(4), 2093–2106. 10.1016/S0006-3495(00)76756-7

43. Koonin, E. V., Wolf, Y. I., & Karev, G. P. (2002). The structure of the protein universe and genome evolution. Nature 2002 420:6912, 420(6912), 218–223. 10.1038/nature01256

44. Larkin, M. A., Blackshields, G., Brown, N. P., Chenna, R., Mcgettigan, P. A., McWilliam, H., Valentin, F., Wallace, I. M., Wilm, A., Lopez, R., Thompson, J. D., Gibson, T. J., & Higgins, D. G. (2007). Clustal W and Clustal X version 2.0. Bioinformatics, 23(21), 2947–2948. 10.1093/BIOINFORMATICS/BTM404

45. Lees, J. G., Dawson, N. L., Sillitoe, I., & Orengo, C. A. (2016). Functional innovation from changes in protein domains and their combinations. Current Opinion in Structural Biology, 38, 44–52. 10.1016/J.SBI.2016.05.016

46. Lefort, V., Desper, R., & Gascuel, O. (2015). FastME 2.0: A Comprehensive, Accurate, and Fast Distance-Based Phylogeny Inference Program. Molecular Biology and Evolution, 32(10), 2798–2800. 10.1093/MOLBEV/MSV150

47. Letunic, I., & Bork, P. (2024). Interactive Tree of Life (iTOL) v6: recent updates to the phylogenetic tree display and annotation tool. Nucleic Acids Research, 52(W1), W78–W82. 10.1093/NAR/GKAE268

48. Lopes, T. J. S., Rios, R., Nogueira, T., & Mello, R. F. (2021). Protein residue network analysis reveals fundamental properties of the human coagulation factor VIII. Scientific Reports 2021 11:1, 11(1), 1–11. 10.1038/s41598-021-92201-3

49. Lorch, M., Mason, J. M., Clarke, A. R., & Parker, M. J. (1999). Effects of core mutations on the folding of a β-sheet protein: Implications for backbone organization in the I-state. Biochemistry, 38(4), 1377–1385. 10.1021/BI9817820/ASSET/IMAGES/LARGE/BI9817820FD02A.JPEG

50. Lorch, M., Mason, J. M., Sessions, R. B., & Clarke, A. R. (2000). Effects of mutations on the thermodynamics of a protein folding reaction: Implications for the mechanism of formation of the intermediate and transition states. Biochemistry, 39(12), 3480–3485. 10.1021/BI9923510/SUPPL_FILE/BI9923510_S.PDF

51. Margelevičius, M., & Venclovas, Č. (2010). Detection of distant evolutionary relationships between protein families using theory of sequence profile-profile comparison. BMC Bioinformatics, 11. 10.1186/1471-2105-11-89

52. McGuffin, L. J., Bryson, K., & Jones, D. T. (2001). What are the baselines for protein fold recognition? Bioinformatics, 17(1), 63–72. 10.1093/BIOINFORMATICS/17.1.63

53. Moréra, S., Vigouroux, A., & Stubbs, K. A. (2011). A potential fortuitous binding of inhibitors of an inverting family GH9 β-glycosidase derived from isofagomine. Organic & Biomolecular Chemistry, 9(17), 5945–5947. 10.1039/C1OB05766A

54. Mutt, E., Mathew, O. K., & Sowdhamini, R. (2014). LenVarDB: database of length-variant protein domains. Nucleic Acids Research, 42(D1), D246–D250. 10.1093/NAR/GKT1014

55. Nacher, J. C., Hayashida, M., & Akutsu, T. (2010). The role of internal duplication in the evolution of multi-domain proteins. Biosystems, 101(2), 127–135. 10.1016/J.BIOSYSTEMS.2010.05.005

56. Naveenkumar, N., Kumar, G., Sowdhamini, R., Srinivasan, N., & Vishwanath, S. (2019). Fold combinations in multi-domain proteins. Bioinformation, 15(5), 342. 10.6026/97320630015342

57. Naveenkumar, N., Sowdhamini, R., & Srinivasan, N. (2019). Specialized structural and functional roles of residues selectively conserved in subfamilies of the pleckstrin homology domain family. FEBS Open Bio, 9(11), 1848–1859. 10.1002/2211-5463.12725

58. Orengo, C. A., Michie, A. D., Jones, S., Jones, D. T., Swindells, M. B., & Thornton, J. M. (1997). CATH – a hierarchic classification of protein domain structures. Structure, 5(8), 1093–1109. 10.1016/S0969-2126(97)00260-8

59. Panchenko, A. R., & Madej, T. (2005). Structural similarity of loops in protein families: Toward the understanding of protein evolution. BMC Evolutionary Biology, 5(1), 1–8. 10.1186/1471-2148-5-10/FIGURES/3

60. Prabantu, V. M., Gadiyaram, V., Vishveshwara, S., & Srinivasan, N. (2022). Understanding structural variability in proteins using protein structural networks. Current Research in Structural Biology, 4, 134–145. 10.1016/J.CRSTBI.2022.04.002

61. Prabantu, V. M., Gadiyaram, V., Vishveshwara, S., & Srinivasan, N. (2023). Comparison of structural networks across homologous proteins. Proteins: Structure, Function, and Bioinformatics. 10.1002/PROT.26650

62. Prabantu, V. M., Naveenkumar, N., & Srinivasan, N. (2021). Influence of Disease-Causing Mutations on Protein Structural Networks. Frontiers in Molecular Biosciences, 7, 620554. 10.3389/FMOLB.2020.620554/BIBTEX

63. Prabantu, V. M., Tandon, H., Sandhya, S., Sowdhamini, R., & Srinivasan, N. (2023). The alteration of structural network upon transient association between proteins studied using graph theory. *Proteins: Structure*, Function, and Bioinformatics. 10.1002/PROT.26606

64. Prada Hernández, J. A., Haire, R. N., Allaire, M., Jakoncic, J., Stojanoff, V., Cannon, J. P., Litman, G. W., & Ostrov, D. A. (2006). Ancient evolutionary origin of diversified variable regions demonstrated by crystal structures of an immune-type receptor in amphioxus. Nature Immunology 2006 7:8, 7(8), 875–882. 10.1038/ni1359

65. Qian, J., Luscombe, N. M., & Gerstein, M. (2001). Protein family and fold occurrence in genomes: power-law behaviour and evolutionary model. Journal of Molecular Biology, 313(4), 673–681. 10.1006/JMBI.2001.5079

66. Reeves, G. A., Dallman, T. J., Redfern, O. C., Akpor, A., & Orengo, C. A. (2006). Structural Diversity of Domain Superfamilies in the CATH Database. Journal of Molecular Biology, 360(3), 725–741. 10.1016/J.JMB.2006.05.035

67. Rodrigues, J. P. G. L. M., Teixeira, J. M. C., Trellet, M., & Bonvin, A. M. J. J. (2018). pdb-tools: a swiss army knife for molecular structures. F1000Research 2018 7:1961, 7, 1961. 10.12688/f1000research.17456.1

68. Roy, A., Srinivasan, N., & Gowri, V. S. (2009). Molecular and Structural Basis of Drift in the Functions of Closely-Related Homologous Enzyme Domains: Implications for Function Annotation Based on Homology Searches and Structural Genomics. In Silico Biology, 9(1–2), 41–55. 10.3233/ISB-2009-0379

69. Sandhya, S., Rani, S. S., Pankaj, B., Govind, M. K., Offmann, B., Srinivasan, N., & Sowdhamini, R. (2009). Length Variations amongst Protein Domain Superfamilies and Consequences on Structure and Function. PLoS ONE, 4(3), e4981. 10.1371/JOURNAL.PONE.0004981

70. Schaefer, C., Schlessinger, A., & Rost, B. (2010). Protein secondary structure appears to be robust under in silico evolution while protein disorder appears not to be. Bioinformatics, 26(5), 625–631. 10.1093/BIOINFORMATICS/BTQ012

71. Scholz, M. (2010). Node similarity as a basic principle behind connectivity in complex networks. Journal of Data Mining & Digital Humanities, 2015. 10.46298/jdmdh.33

72. Schubot, F. D., Kataeva, I. A., Chang, J., Shah, A. K., Ljungdahl, L. G., Rose, J. P., & Wang, B.-C. (2004). Structural Basis for the Exocellulase Activity of the Cellobiohydrolase CbhA from Clostridium thermocellum †. 10.1021/bi030202i

73. Seera, S., & Nagarajaram, H. A. (2021). Effect of Disease Causing Missense Mutations on Intrinsically Disordered Regions in Proteins. Protein & Peptide Letters, 29(3), 254–267. 10.2174/0929866528666211126161200

74. Sen, N., & Madhusudhan, M. S. (2022). A structural database of chain–chain and domain–domain interfaces of proteins. Protein Science, 31(9), e4406. 10.1002/PRO.4406

75. Siltberg-Liberles, J., Grahnen, J. A., & Liberles, D. A. (2011). The Evolution of Protein Structures and Structural Ensembles Under Functional Constraint. Genes, 2(4), 748. 10.3390/GENES2040748

76. S.J. Hubbard, & J.M. Thornton. (1993). NACCESS. Http://Www.Bioinf.Manchester.Ac.Uk/Naccess/.

77. Tan, K., Chang, C., Cuff, M., Osipiuk, J., Landorf, E., Mack, J. C., Zerbs, S., Joachimiak, A., & Collart, F. R. (2013). Structural and functional characterization of solute binding proteins for aromatic compounds derived from lignin: p-Coumaric acid and related aromatic acids. Proteins: Structure, Function, and Bioinformatics, 81(10), 1709–1726. 10.1002/PROT.24305

78. Taverna, D. M., & Goldstein, R. A. (2002). Why are proteins so robust to site mutations? Journal of Molecular Biology, 315(3), 479–484. 10.1006/JMBI.2001.5226

79. Teichmann, S. A., Park, J., & Chothia, C. (1998). Structural assignments to the Mycoplasma genitalium proteins show extensive gene duplications and domain rearrangements. 95, 14658–14663. www.tigr.org.

80. Thangudu, R. R., Manoharan, M., Srinivasan, N., Cadet, F., Sowdhamini, R., & Offmann, B. (2008). Analysis on conservation of disulphide bonds and their structural features in homologous protein domain families. BMC Structural Biology, 8(1), 1–22. 10.1186/1472-6807-8-55/TABLES/2

81. Todd, A. E., Orengo, C. A., & Thornton, J. M. (2001). Evolution of function in protein superfamilies, from a structural perspective. Journal of Molecular Biology, 307(4), 1113–1143. 10.1006/JMBI.2001.4513

82. Tokuriki, N., & Tawfik, D. S. (2009). Stability effects of mutations and protein evolvability. Current Opinion in Structural Biology, 19(5), 596–604. 10.1016/J.SBI.2009.08.003

83. Trakhanov, S., Vyas, N. K., Luecke, H., Kristensen, D. M., Ma, J., & Quiocho, F. A. (2005). Ligand-free and -bound structures of the binding protein (LivJ) of the Escherichia coli ABC leucine/isoleucine/valine transport system: Trajectory and dynamics of the interdomain rotation and ligand specificity. Biochemistry, 44(17), 6597–6608. 10.1021/BI047302O/ASSET/IMAGES/LARGE/BI047302OF00005.JPEG

84. Vijayabaskar, M. S., & Vishveshwara, S. (2012). Insights into the Fold Organization of TIM Barrel from Interaction Energy Based Structure Networks. PLOS Computational Biology, 8(5), e1002505. 10.1371/JOURNAL.PCBI.1002505

85. Vogel, C., Bashton, M., Kerrison, N. D., Chothia, C., & Teichmann, S. A. (2004). Structure, function and evolution of multidomain proteins. Current Opinion in Structural Biology, 14(2), 208–216. 10.1016/J.SBI.2004.03.011

86. Vogel, C., Berzuini, C., Bashton, M., Gough, J., & Teichmann, S. A. (2004). Supra-domains: Evolutionary Units Larger than Single Protein Domains. Journal of Molecular Biology, 336(3), 809–823. 10.1016/j.jmb.2003.12.026

87. Vogel, C., Teichmann, S. A., & Pereira-Leal, J. (2005). The Relationship Between Domain Duplication and Recombination. Journal of Molecular Biology, 346(1), 355–365. 10.1016/J.JMB.2004.11.050

88. Wang, G., & Dunbrack, R. L. (2003). PISCES: a protein sequence culling server. Bioinformatics (Oxford, England), 19(12), 1589–1591. 10.1093/BIOINFORMATICS/BTG224

89. Wang, N., Du, N., Peng, Y., Yang, K., Shu, Z., Chang, K., Wu, D., Yu, J., Jia, C., Zhou, Y., Li, X., Liu, B., Gao, Z., Zhang, R., & Zhou, X. (2021). Network Patterns of Herbal Combinations in Traditional Chinese Clinical Prescriptions. Frontiers in Pharmacology, 11, 590824. 10.3389/FPHAR.2020.590824/FULL

90. Zhang, Y., & Skolnick, J. (2005). TM-align: a protein structure alignment algorithm based on the TM-score. Nucleic Acids Research, 33(7), 2302–2309. 10.1093/NAR/GKI524

91. Zhu, H., Domingues, F. S., Sommer, I., & Lengauer, T. (2006). NOXclass: prediction of protein-protein interaction types. BMC Bioinformatics, 7(1), 27. 10.1186/1471-2105-7-27

