## Supplementary Material for "Structural network comparison of domains across remote homologues arising from different domain architectures"

### Supplementary documents

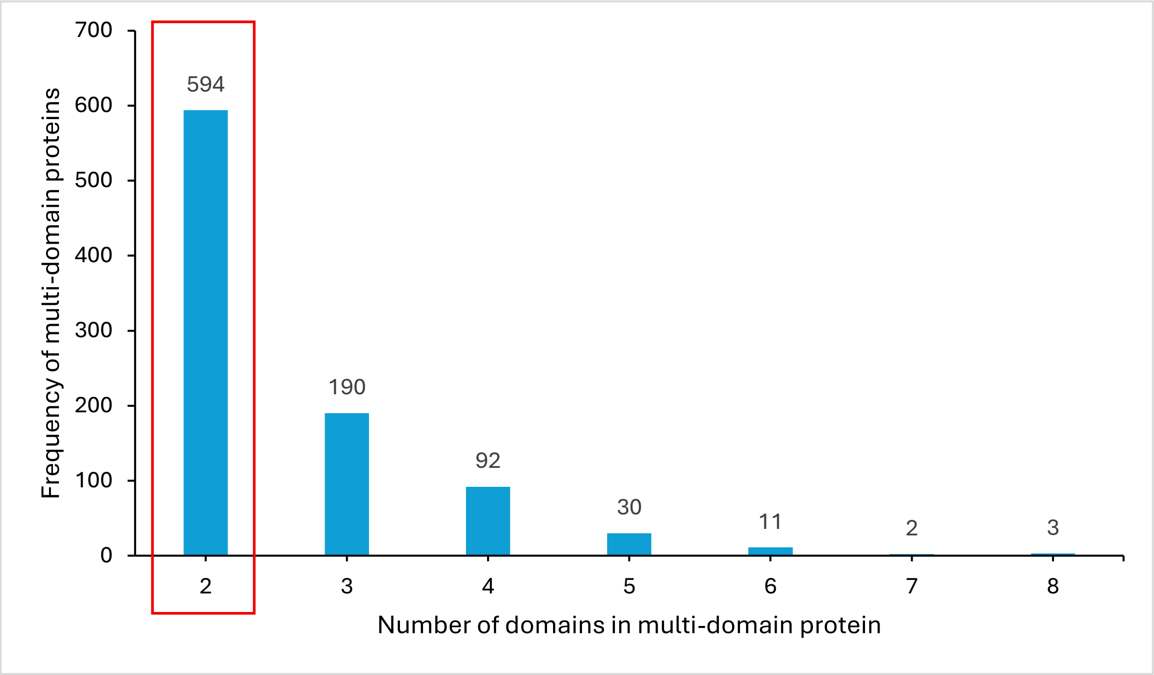

**Figure S1 Frequency of multi-domain proteins in the initial dataset**. Only two-domain proteins were considered in this study and highlighted with a red outline in the figure.

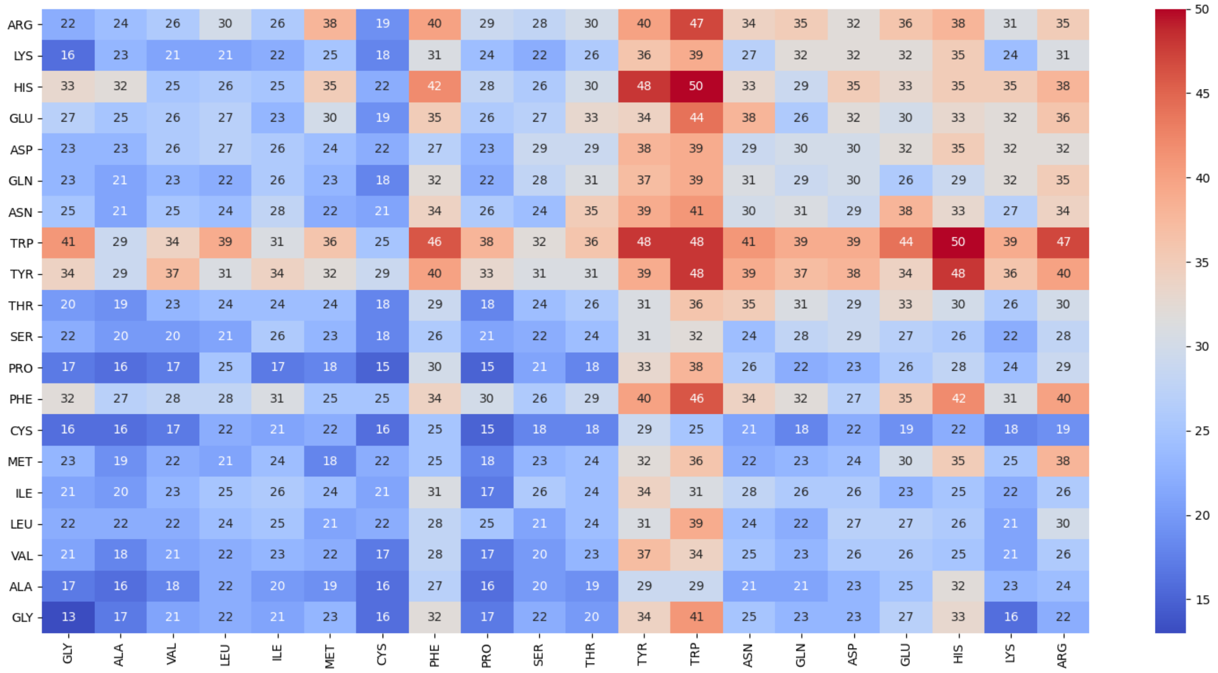

**Figure S2 Maximum number of atomic contacts per amino acid residue pairs in the whole dataset.** The cells are marked with the maximum number of observed atomic contacts and coloured accordingly.

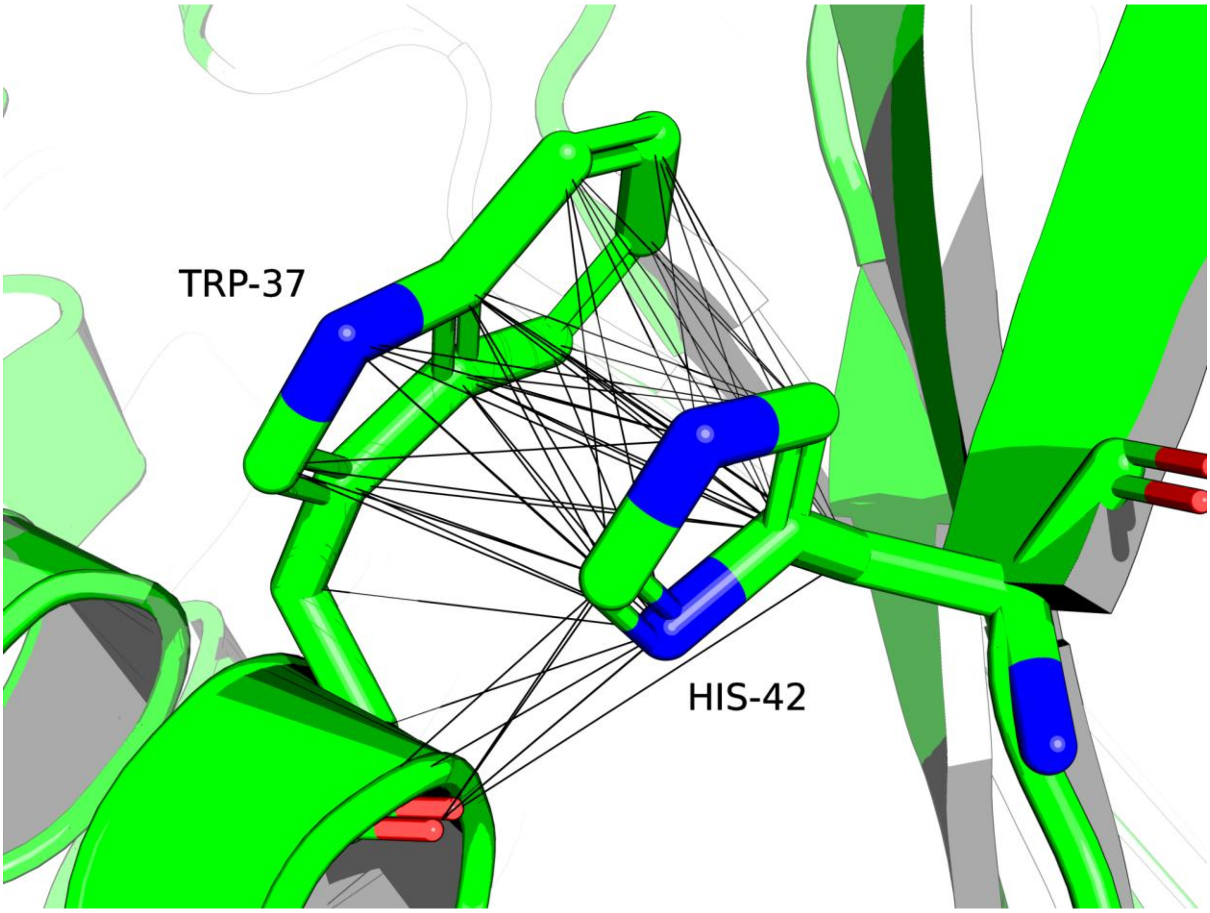

**Figure S3 Illustration of atomic contacts.** The largest number of atomic contacts (~50 contacts) formed by Tryptophan and Histidine residues from the second domain of Phosphoglycerate mutase (PDB ID: 1EQJ)

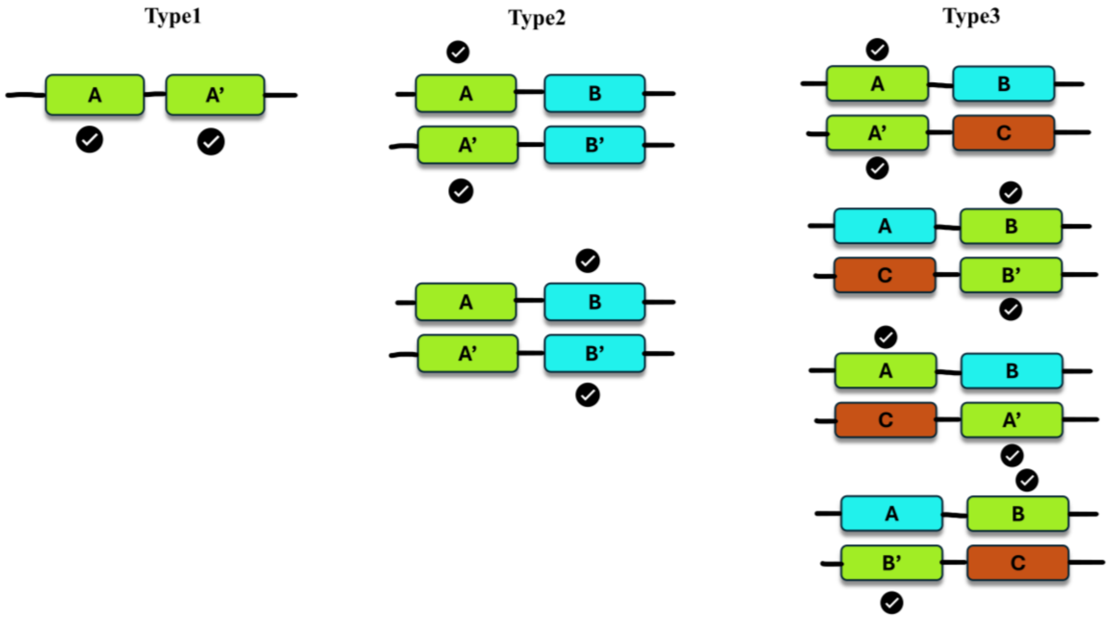

**Figure S4 All possible domain pairs used in the study**. The pairs considered for comparison are marked with tick mark.

**Table S1 Number of peculiar pairs in Type 2 and Type 3**. The subtypes correspond to all possible types in supplementary figure S4 as shown in order. Protein1 or protein2 correspond to number of proteins having Type 1 like architecture from which domains are taken for comparison. Total number of peculiar pairs in Type 2 will be same as protein1 or protein2 as the rule makes them comparisons of domains from two Type1 pairs.

| Types | Protein1 | Protein2 | Total (#Peculiar/#Total) | Total | |
|---|---|---|---|---|---|
| Type2_1 | 189 | 189 | 189/414 | 391/834 | Type2 |
| Type2_2 | 202 | 202 | 202/420 | | |
| Type3_1 | 4 | 8 | 12/139 | 53/270 | Type3 |
| Type3_2 | 0 | 9 | 9/25 | | |
| Type3_3 | 14 | 2 | 16/52 | | |
| Type3_4 | 2 | 14 | 16/54 | | |

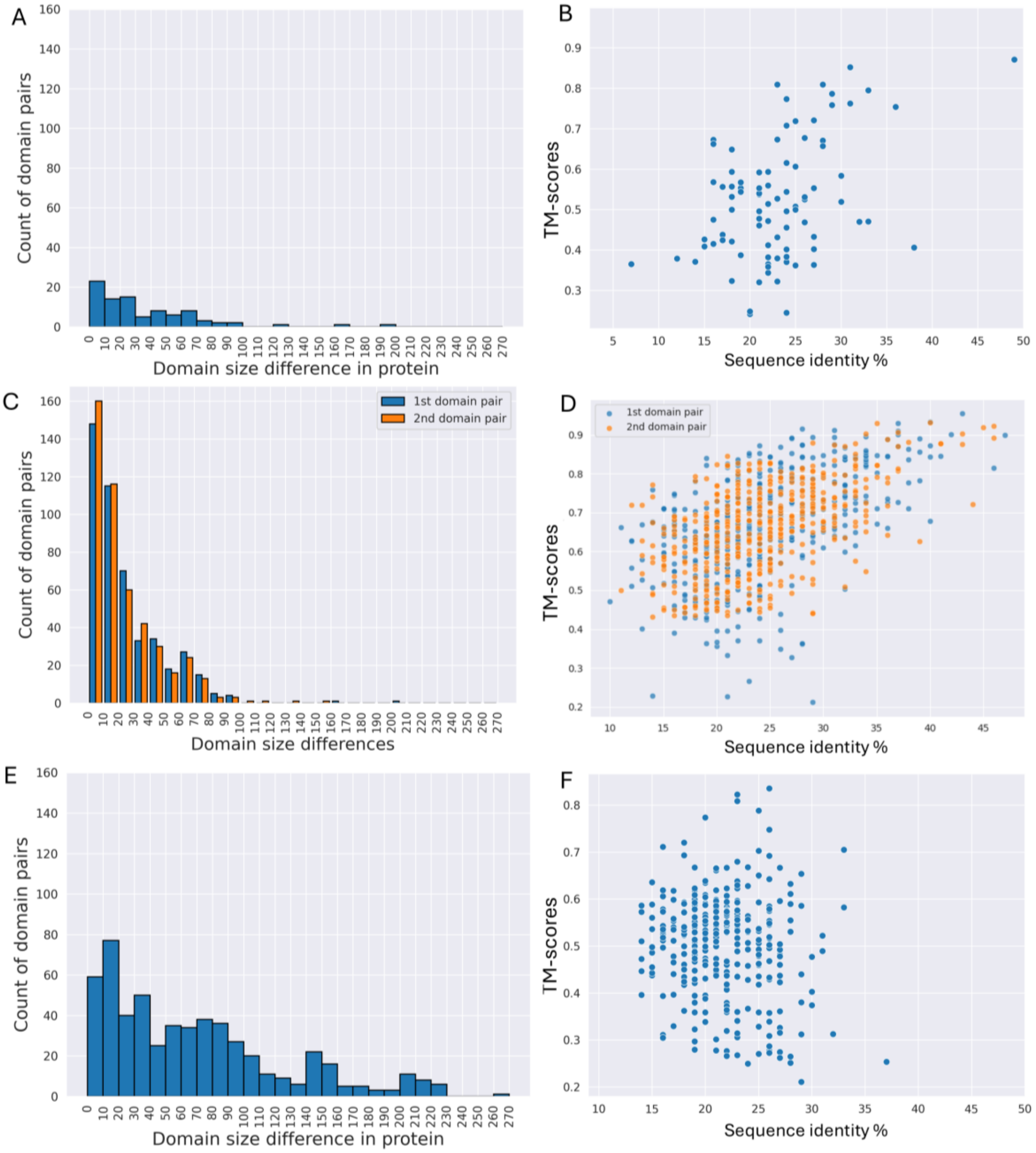

**Figure S5 Preliminary domain comparisons**. The left panel consisting of sub-figures A, C and E shows domain size differences based on sequence length and the frequencies. The right panel consisting of sub-figures B, D and F shows the sequence identities in the x-axis and TM scores in the y-axis. For type2 (sub-figure C and D) both N and C-terminal pairs are shown together.

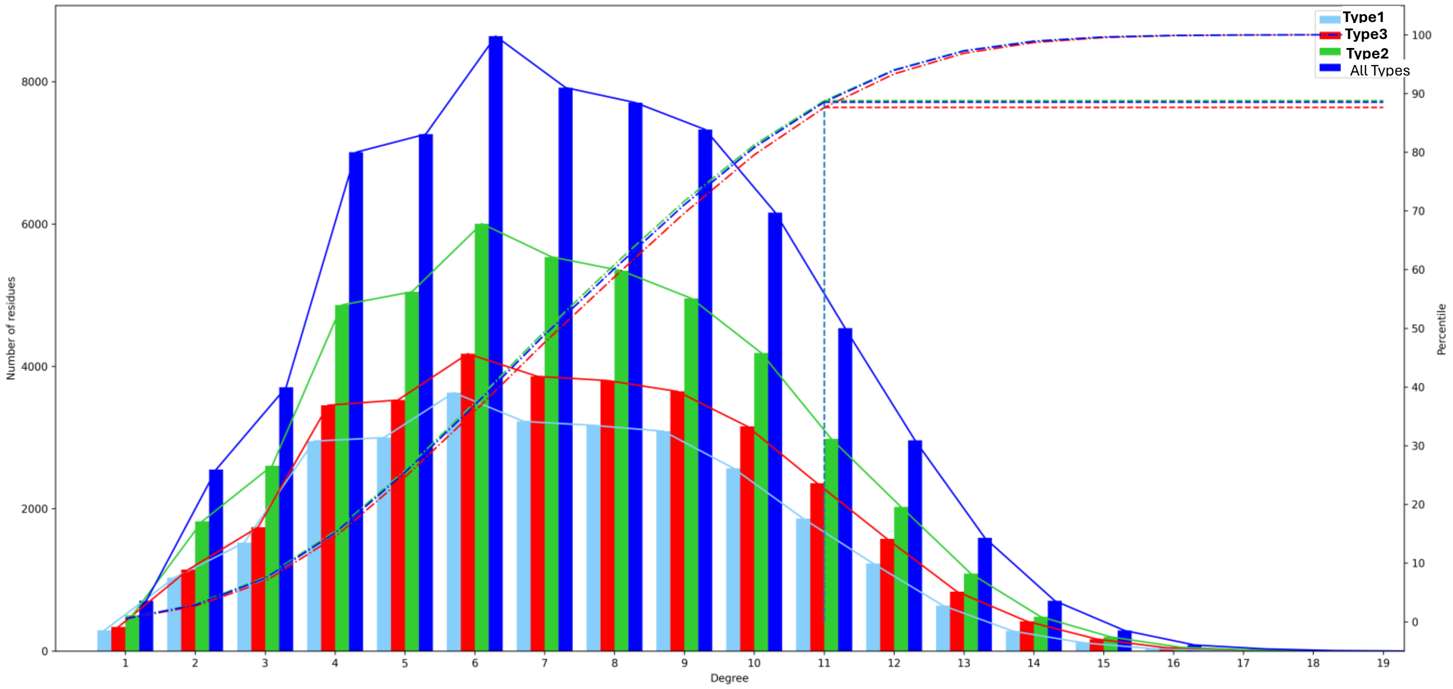

**Figure S6 Degree distribution of nodes in the dataset**. Distribution of different domain pairs are shown in different colours. The secondary y-axis shows the percentile score. The dash-dot lines show the percentiles of each degree. The dashed line points the degree corresponding to the 90$^{th}$ percentile.

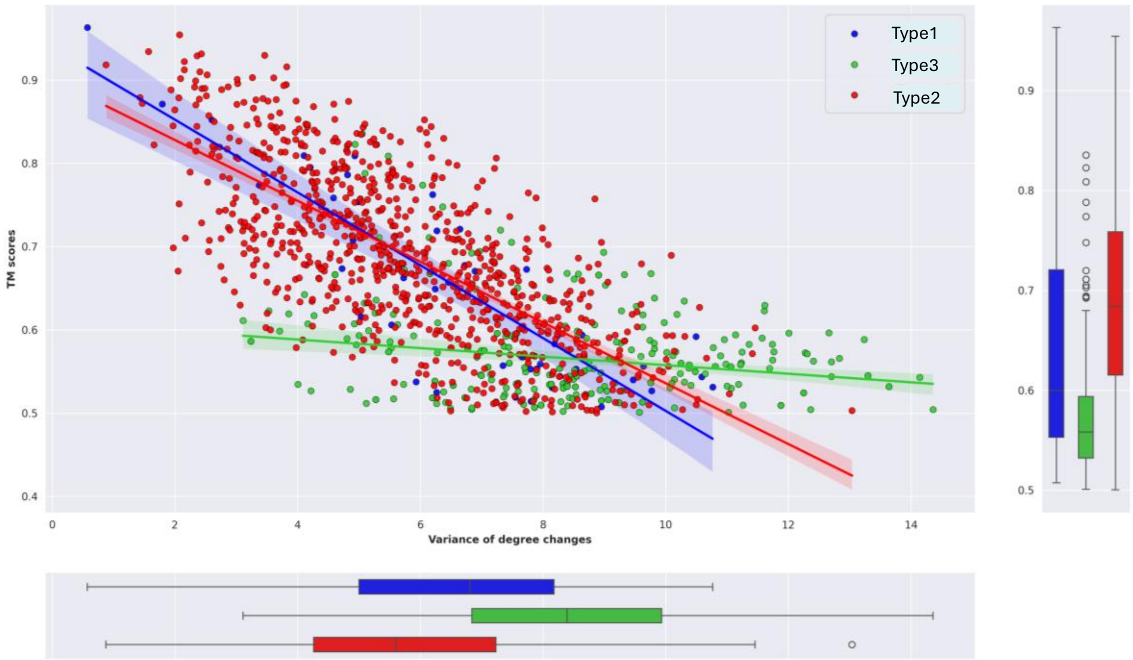

**Figure S7 Relationship between degree difference variances (degree variability) and TM-scores**. The distribution of these variables is shown in a marginal plot. The lines in the scatterplot are the linear regression lines.

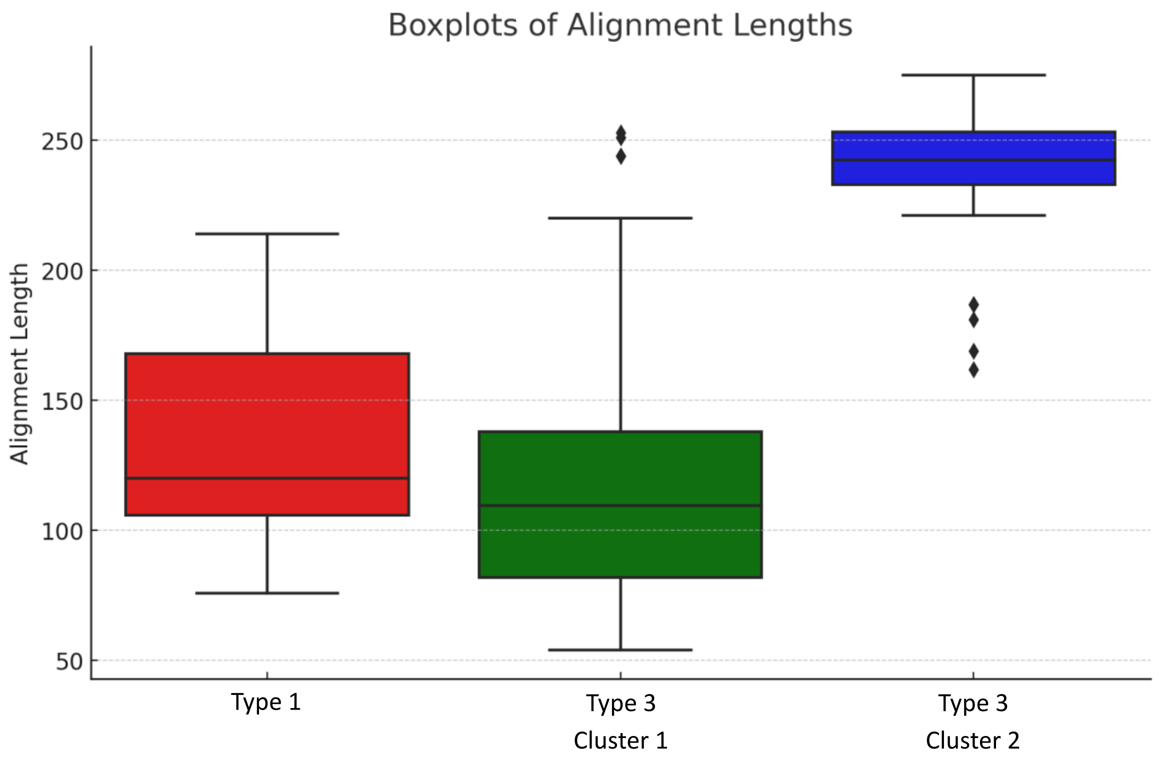

**Figure S8 Alignment lengths of domain pairs in Type1 and Type3**. Distribution of alignment lengths of Type 1 and Type 3. Type 3 is divided into two clusters. Conserved hubs of equal to or less than 30 and non-conserved hubs equal to or less than 50 comprises cluster 1. Cluster 2 spans non-conserved hubs greater than 50.

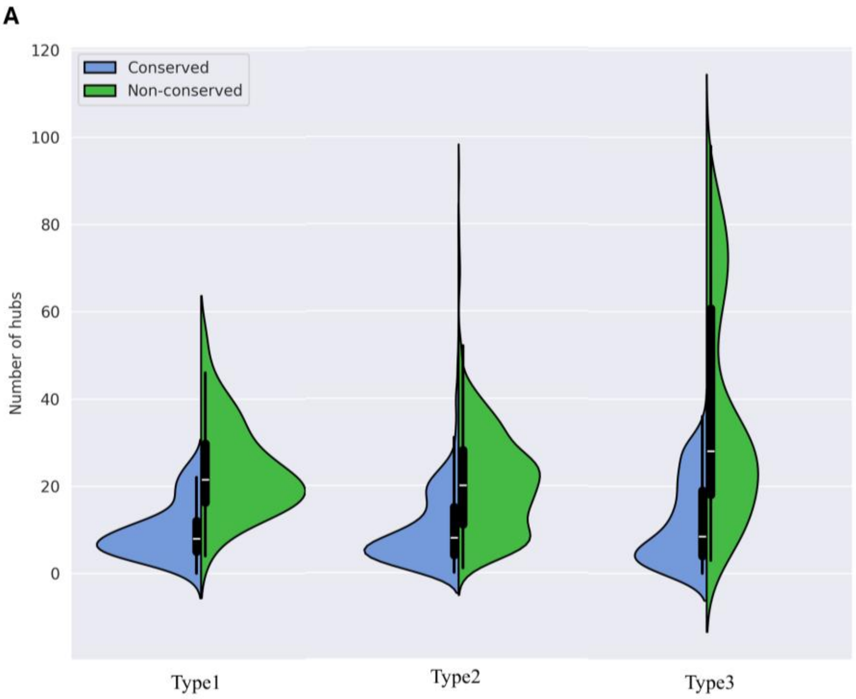

**B**

| | Type1 | Type2 | Type3 |
|---|---|---|---|
| Type1 | - | 0.029 (*) | 0.004 (**) |
| Type2 | 0.337 (NS) | - | $1.35*10^{-19}$ (****) |
| Type3 | 0.302 (NS) | 0.052 (NS) | - |

**Figure S9 Hub preservation in domain pairs**. **A.** Distribution of both conserved and non-conserved hubs are shown in spit violin plot. **B.** Mann Whitney U p-Values are mentioned in a table where lower triangle shows value for distribution of conserved hubs among types while upper triangle shows value for distribution of non-conserved hubs.

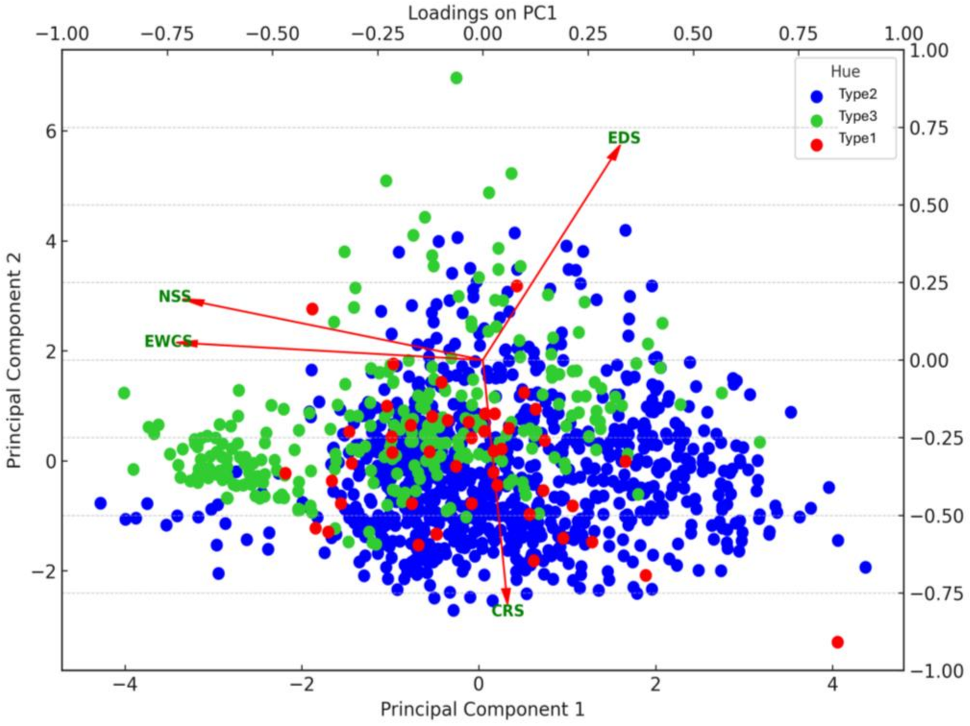

**Figure S10 Decomposition of Network similarity (NSS) and its components into principal components**. The secondary axis shows the loadings of each feature on PCs.

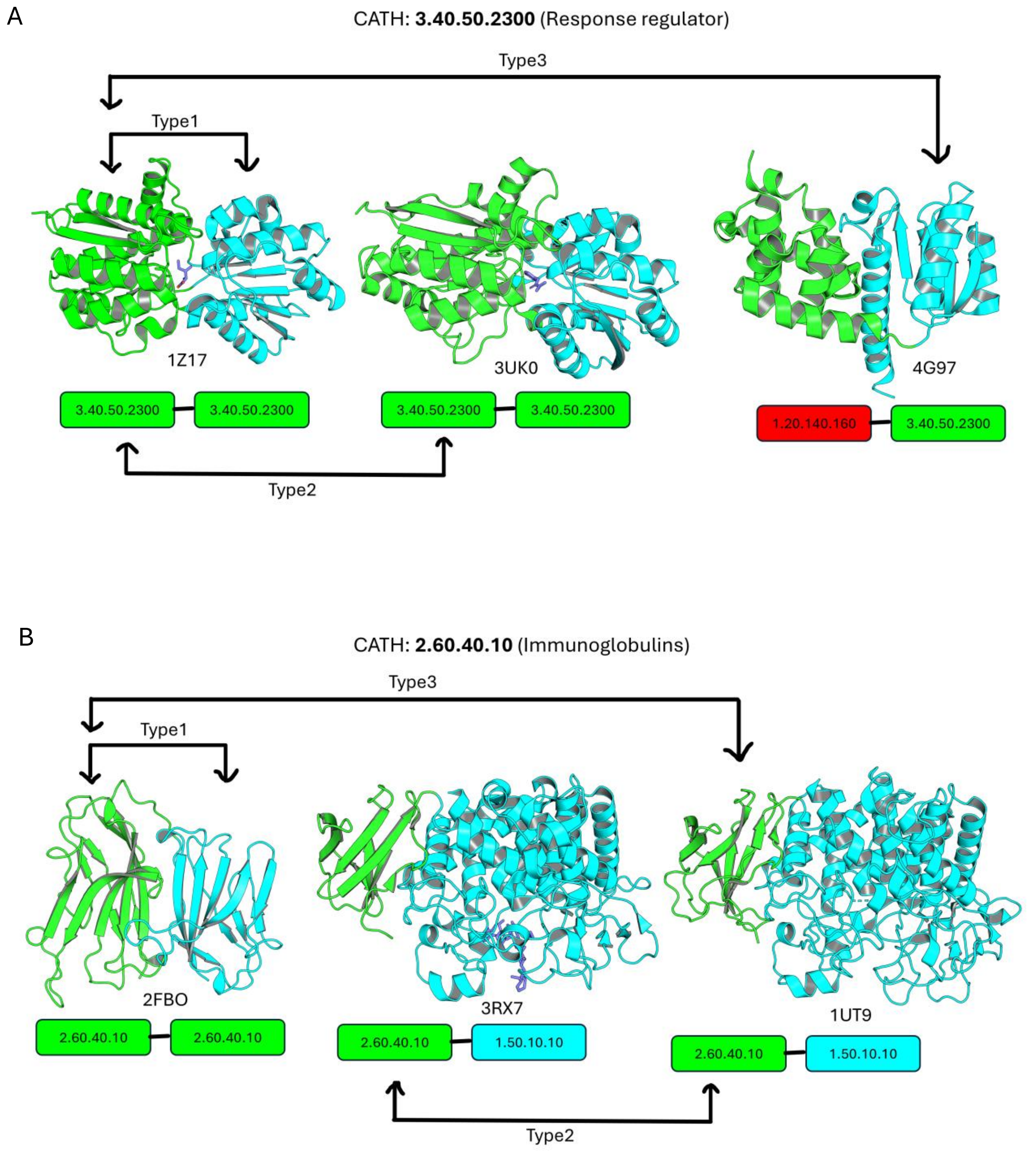

**Figure S11 Domain combinations in case studies**. Two domains in each cartoon structure are shown in green (dom1) and cyan (dom2) and types of domain comparisons are labelled with arrows. CATH ids of each protein are shown below the cartoon structure in boxes, where a different interacting domain partner in proteins is coloured other than green (like cyan and red) similar to supplementary figure 1. **A**. Homologous superfamily Response regulator and **B.** Immunoglobulins. In some structures, a ligand is shown in the interdomain cleft in blue colour.

**Table S2 Properties of amino acid residues showing high degree differences** in superfamily **2.60.40.10 (Immunoglobulins)**. #DD represents the difference of degrees between residue pairs. The degrees are substracted from left to right and a negative sign indicates the right side residue with a higher degree. The properties to look for is shown in bold and satisfies our hypothesis. In some instances it is shown in both bold and italic font when a relaxed solvent accessibility cutoff is used to show buridness. **RSA**: Relative solvent accessibility. The percentage change in number of atoms was calculated by (max(no. of atoms)-lowest(no. of atoms))*100/max(no. of atoms). Wherever the change in atom number exceeds 40%, it is shown in bold. For clustering coefficients, if the difference is around 0.1, it is shown in bold font.

| Type1 (2FBO) | | | | |
|---|---|---|---|---|
| **Residues** | **#DD** | **RSA** | **Δ atoms (%change)** | **Clustering coefficient** |
| K43 <--> T174 | 6 | 20.6 <--> 25.3 | 9 <--> 7 (22) | 0.19 <--> 0.20 |
| F54 <--> A191 | 6 | 1.9 <--> 0.1 | 11 <--> 5 (**54**) | **0.18** <--> **0.32** |
| L89 <--> A214 | 10 | **0.1** <--> **56.2** | 8 <--> 5 (37.5) | **0.21** <--> **0.33** |
| Q113 <--> T236 | 6 | ***14.5*** <--> ***51.6*** | 9 <--> 7 (22) | **0.31** <--> **0.5** |
| Type2 (3RX7_1UT9) | | | | |
| S8 <--> D213 | -6 | ***34.3*** <--> ***10.5*** | 6 <--> 8 (25) | **0.40** <--> **0.23** |
| N55 <--> Y268 | -6 | 79.5 <--> 30.4 | 8 <--> 12 (33) | **0.30** <--> **0.14** |
| Type3 (2FBO_1UT9) | | | | |
| M5 <--> P212 | 6 | 4.2 <--> 2.3 | 8 <--> 7 (12.5) | **0.29** <--> **0.38** |
| V22 <--> G225 | 7 | **4.8** <--> **57.9** | 7 <--> 4 (**42.8**) | **0.31** <--> **0.66** |
| P54 <--> E253 | 10 | **1.9** <--> **58.8** | 7 <--> 9 (22.2) | **0.18** <--> **0.33** |
| A90 <--> H270 | -9 | **49.6** <--> **0.0** | 5 <--> 10 (**50**) | **0.5** <--> **0.23** |
| L96 <--> D276 | 10 | **0.2** <--> **103.6** | 8 <--> 8 (0) | 0.34 <--> 0 |
| V99 <--> A278 | 9 | **1.4** <--> **33.3** | 7 <--> 5 (28.5) | 0.29 <--> 0.3 |
| D103 <--> T279 | 7 | **3.2** <--> **58** | 8 <--> 7 (12.5) | **0.36** <--> **0.83** |
| Q113 <--> T290 | 6 | ***14.5*** <--> ***69.8*** | 9 <-->7 (22.2) | 0.31 <--> 0.33 |

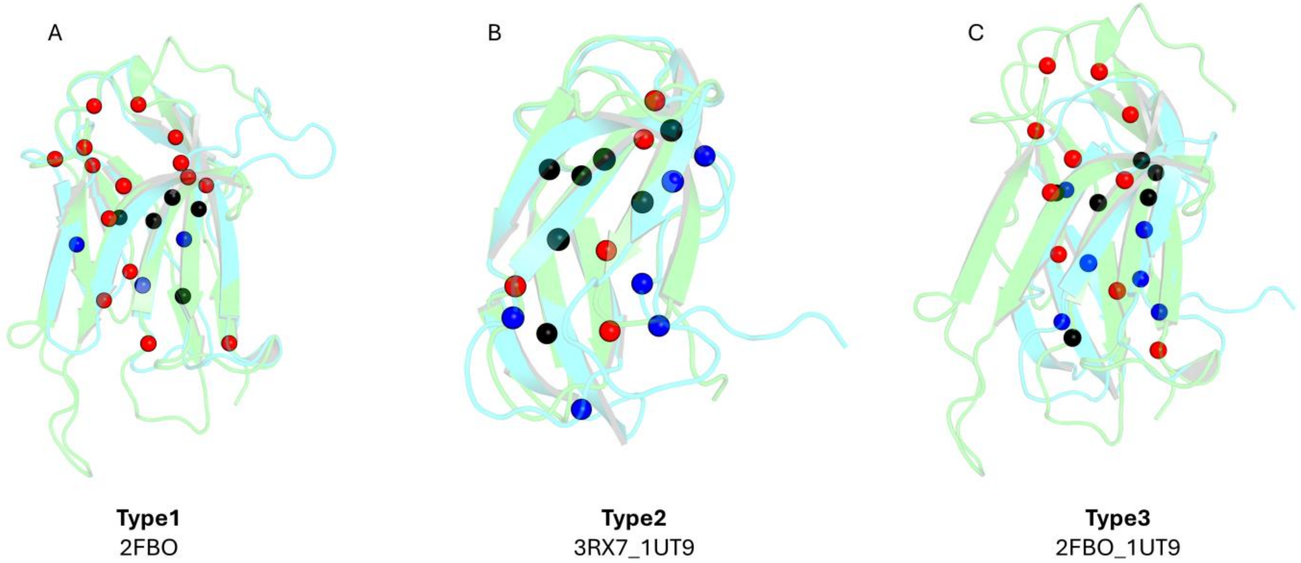

**Figure S12 Distribution of hubs** in superfamily **Immunoglobulins (CATH: 2.60.40.10)**. Homologous superfamily domains in types of comparison, shown in A, B, C are superposed where one domain is coloured green and the other cyan. Black spheres show conserved hubs between the two networks. Red spheres show unique hubs to domain1 (green cartoon) and blue spheres show unique hubs to domain2 (cyan cartoon).

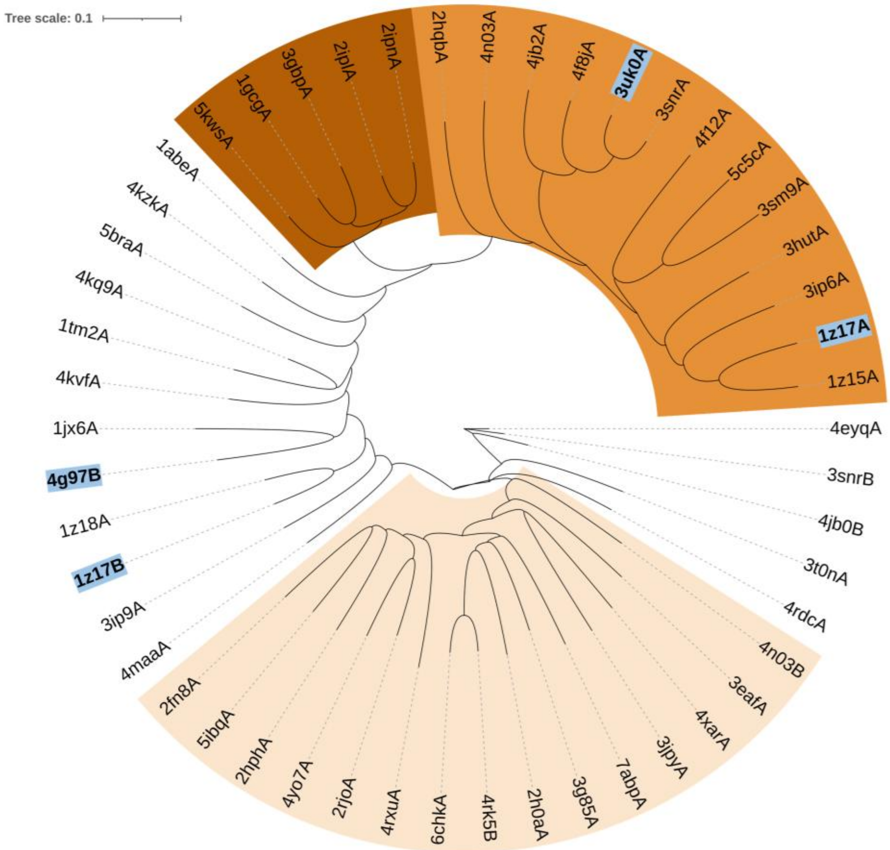

**Figure S13 Phylogenetic dendrogram of Network dissimilarities** of homologous superfamily domains from Response regulators. The case studies are shown in bold font and light blue background. Clades are coloured based on visual clade formation.

**Table S3 Network parameters of the case studies.** The differences between the parameters are shown withing brackets.

| | CATH: 3.40.50.2300 (Response regulator) | | | CATH:2.60.40.10 (Immunoglobulins) | | |
|---|---|---|---|---|---|---|
| **Network property** | **Type1 (1Z17)** | **Type2 (1Z17_3UK0)** | **Type3 (1Z17_4G97)** | **Type1 (2FBO)** | **Type2 (3RX7_1UT9)** | **Type3 (2FBO_1UT9)** |
| Degree variance | 8.27 | 6.45 | 9.27 | 4 | 2 | 8 |
| NSS | 0.359 | 0.35 | 0.373 | 0.3 | 0.208 | 0.377 |
| RMSD | 2.98 | 2.22 | 3.33 | 1.99 | 1.62 | 3.08 |
| TM score | 0.544 | 0.8 | 0.526 | 0.762 | 0.768 | 0.5 |
| Sequence identity % | 24 | 22 | 21 | 31 | 32 | 20 |
| Conserved/non-conserved hubs | 12/32 | 19/35 | 7/30 | 5/18 | 7/11 | 6/16 |
| Unique hubs | 17 vs 15 | 21 vs 14 | 26 vs 4 | 15 vs 3 | 5 vs 6 | 10 vs 6 |
| Mean clustering coefficient | 0.298 vs 0.313 (0.015) | 0.298 vs 0.308 (0.01) | 0.298 vs 0.276 (0.022) | 0.302 vs 0.320 (0.018) | 0.290 vs 0.327 (0.037) | 0.302 vs 0.327 (0.025) |
| Network density | 0.039 vs 0.053 (0.014) | 0.039 vs 0.038 (0.01) | 0.039 vs 0.053 (0.014) | 0.05 vs 0.056 (0.006) | 0.079 vs 0.066 (0.013) | 0.05 vs 0.066 (0.016) |
| Avg. shortest path length | 3.878 vs 3.459 (0.419) | 3.878 vs 4.014 (0.136) | 3.878 vs 3.696 (0.18) | 3.787 vs 3.654 (0.133) | 3.071 vs 3.340 (0.269) | 3.787 vs 3.340 (0.447) |
| Degree assortativity | 0.259 vs 0.332 (0.073) | 0.259 vs 0.110 (0.149) | 0.259 vs 0.208 (0.051) | 0.198 vs 0.206 (0.008) | 0.135 vs 0.102 (0.033) | 0.198 vs 0.102 (0.004) |
| Network diameter | 8 vs 7 (1) | 8 vs 9 (1) | 8 vs 9 (1) | 10 vs 8 (2) | 7 vs 8 (1) | 10 vs 8 (2) |
| Transitivity | 0.274 vs 0.268 (0.006) | 0.274 vs 0.267 (0.007) | 0.274 vs 0.261 (0.013) | 0.282 vs 0.263 (0.019) | 0.285 vs 0.266 (0.019) | 0.282 vs 0.266 (0.016) |
| Betweenness MSE | 0.0004 | 0.0001 | 0.0005 | 0.0003 | 0.0009 | 0.0006 |
| Closeness MSE | 0.001 | 0.0003 | 0.008 | 0.0004 | 0.006 | 0.0018 |

**Table S4 Mapping of functional sites in domain comparisons**. The residues are outlined an oval shape if it is a functionally important residue as mentioned in the literature. If the residues are hubs in our network study, they are additionally outlined with a green box. For type3 domain comparison, the interfacial residues on domain 2 of 4G97 and their topological equivalent residues in 1Z17 are shown in Italics. The residues whose topological equivalence is not found are marked with X.

**Type1 (1Z17)**

| Dom1 | Dom2 | #DD |
|---|---|---|
| A100 | G227 | -2 |
| T102 | A229 | 1 |
| X | E226 | |
| C78 | Y202 | -2 |
| S79 | H203 | -1 |
| Y18 | Y150 | -3 |
| L77 | G201 | 7 |
| A101 | V228 | -1 |
| F276 | X | |
| X | D123 | |
| M313 | F329 | 3 |

**Type2 (1Z17_3UK0)**

| Dom1 | Dom2 | #DD |
|---|---|---|
| A100 | A123 | -4 |
| T102 | P124 | 1 |
| C78 | V101 | -2 |
| S79 | T102 | 1 |
| Y18 | L46 | 0 |
| L77 | S100 | 6 |
| A101 | X | |
| F276 | F306 | 1 |
| A275 | Q305 | 0 |
| T279 | H309 | -1 |
| T117 | P139 | 2 |
| E22 | E50 | 5 |

**Type3 (1Z17_4G97)**

| Dom1 | Dom2 | #DD |
|---|---|---|
| A10 | E146 | 1 |
| G13 | D147 | -3 |
| L77 | D191 | 6 |
| A101 | T219 | 1 |
| R116 | F235 | 6 |
| S79 | Q193 | 4 |
| X | K239 | |
| F276 | N242 | 5 |
| *P104* | *L225* | -4 |
| *E105* | *L226* | -1 |
| *A108* | *R230* | 0 |
| *G110* | *E232* | -5 |
| *I114* | *P233* | 5 |
| *L115* | *T234* | 8 |
| *G119* | *T239* | -2 |
| *Y281* | *K247* | 4 |
| *S286* | *Q252* | 6 |
